# Importin α2 association with chromatin: Direct DNA binding via a novel DNA binding domain

**DOI:** 10.1101/2020.05.04.075580

**Authors:** Kazuya Jibiki, Takashi S. Kodama, Atsushi Suenaga, Yota Kawase, Noriko Shibazaki, Shin Nomoto, Seiya Nagasawa, Misaki Nagashima, Shieri Shimodan, Renan Kikuchi, Mina Okayasu, Ruka Takashita, Rashid Mehmood, Noriko Saitoh, Yoshihiro Yoneda, Ken-ichi Akagi, Noriko Yasuhara

## Abstract

Nuclear transport of proteins is important for facilitating appropriate nuclear functions. The proteins of the importin α family play key roles in nuclear transport as transport receptors for a huge number of nuclear proteins. Additionally, these proteins possess other functions, including chromatin association and gene regulation. However, these non-transport functions of importin α are not yet fully understood, especially their molecular-level mechanisms for functioning with chromatin and their consequences. Here, we report the novel molecular characteristics of importin α involving binding to diverse sequences in chromatin. We newly identified and characterized a DNA-binding domain—the Nucleic Acid Associating Trolley pole domain (NAAT domain)—in the N-terminal region of importin α within the conventional importin β binding (IBB) domain, which was shown to be necessary for nuclear transport of cargo proteins. We propose a ‘stroll and locate’ model to explain the association of importin α with chromatin. This is the first study to delineate the interaction between importin α and chromatin DNA via the NAAT domain, indicating the bifunctionality of the importin α N-terminal region for nuclear transport and chromatin association.

## Introduction

The importin α family is a class of nuclear transport receptors that mediate protein translocation from the cytoplasm into the cell nucleus through the nuclear pore in eukaryotic cells (1). Proteins are generally synthesised in the cytoplasm, so nuclear proteins, such as transcription factors, have to be transported into the nucleus via transport receptors such as importins. Importin α receptors recognise their cargo proteins by their nuclear localisation signal (NLS) (2–3). Importin α carries out the nuclear import process by forming a trimeric complex with importin β1 and the cargo protein (4–5). The binding of Ran-GTP to importin β1 dissociates importin β1 from the importin α-cargo protein complex (6), while the binding of Nup50 or CAS (cellular apoptosis susceptibility protein) to importin α facilitates the dissociation of importin α from its cargo (6–8).

Architecturally, importin α family proteins consist of three domains: 1) the N-terminal importin β–binding (IBB) domain, which interacts with importin β1 or otherwise binds in an autoregulatory fashion to two NLS-binding sites of importin α itself (9–10); 2) a main body with a helix repeat domain called armadillo (ARM) repeats, which includes two NLS-binding sites; and 3) the C-terminal region, which includes the Nup50 and the CAS binding domain necessary for NLS release and importin α nuclear export for recycling.

Importin α family proteins are expressed from several gene families in mammalian cells, and these families possess distinct cargo specificities. Their expression profiles vary widely depending on the cell types, and the protein activities in the nucleus are regulated through selective nuclear protein transport (11–13). In this study, we designate the importin α family proteins as importin α1 (KPNA1, NPI1, importin α5 in humans), importin α2 (KPNA2, Rch1, and importin α1 in humans), importin α3 (KPNA3, Qip2, and importin α4 in humans), importin α4 (KPNA4, Qip1, and importin α3 in humans), importin α6 (KPNA6, NPI2, and importin α7 in humans), and importin α8 (KPNA7).

Importin α proteins have been shown to perform other functions in addition to their NLS transport receptor function for selective nuclear transport. These non-canonical functions include spindle assembly, lamin polymerisation, nuclear envelope formation, protein degradation, stress response, gene expression, cell surface function, and mRNA-related functions (11). In addition, importin α family members are also known to accumulate in the nucleus under certain stress conditions, such as heat shock and oxidative stress, wherein they bind to a DNase-sensitive nuclear component (14–17). Thus, for a deeper and more comprehensive understanding of the roles of importin α proteins in cellular events, an overall view of their canonical and non-canonical functions and a thorough elucidation of the underlying molecular mechanisms of each function are important.

Among these non-canonical functions, the association of importin α proteins, especially importin α2, with chromatin was the focus of this study. In the present study, we attempted to investigate the molecular mechanisms underlying this chromatin association of importin α and revealed that importin α proteins directly bind to multiple regions in genomic DNA through a novel chromatin-associating domain in the IBB domain. We also found that the association of importin α2 with DNA was multi-modal, electrostatic, of intermediate strength, and semi-specific. These characteristics of the association allowed importin α to move around the DNA and facilitate efficient delivery of proteins to their target sites and recruit proteins through a facilitated diffusion mechanism, i.e. a ‘stroll and locate’ model. This is the first study to reveal that importin α is a DNA-binding protein with a novel DNA-binding domain essential for the non-canonical role of the IBB domain.

## Materials and methods

### Cell culture

Mouse ES cell lines were cultured as follows. The Bruce-4 cell line was maintained in DMEM supplemented with 15% FCS and ESGRO (Merck Millipore, UK) on a 0.1% gelatine-coated dish at 10% CO_2_.

### Immunostaining

For immunostaining, cells were seeded on 0.1% gelatine-coated cover glass slips and then fixed in 3.7% formalin (Nacalai Tesque Inc., Japan., Japan.) in PBS. Cells were permeabilized using 0.5% Triton X100 (Nacalai Tesque Inc., Japan.) in PBS and blocked with 3% skim milk (Nacalai Tesque Inc., Japan.) in PBS. The primary antibody for KPNA2 (importin α2) (rat monoclonal antibody, MBL, Japan: 1/400 dilution; or goat polyclonal, Santa Cruz: 1/200 dilution) and the secondary antibody (Alexa488 conjugated anti-rat or anti-rabbit IgG in 1/100 dilution; Thermo Fisher Scientific, US) were suspended in Can get signal solution (Toyobo, Japan). DNA was stained with DAPI (Nacalai Tesque Inc., Japan.), and images were obtained by confocal microscopy A1 (Nikon, Japan) equipped with 60X objective lens using the software NIS-Elements (Nikon, Japan).

### Plasmid construction

The full-length wild-type and mutant pEGFP constructs of importin α were first cloned as previously described (17). pEGFP-importin α2 NAAT mutants 28A4, 39A5, and 49A3 were constructed using the KOD-plus-Mutagenesis Kit (Toyobo, Japan). The C-mutants of wild-type and 28A4 importin α2 were constructed according to the previous report (17).

The primers used were as follows:

NAAT-28A4 Fwd-GCTGCTATAGAAGTTAATGTGGAACTCAGGAAA
NAAT-28A4 Rev-AGCAGCCATTTCTGTGCTGTCCTTCCC
NAAT-39A5 Fwd-AGCGGCAGCGAGTTCCACATTAAC
NAAT-39A5 Rev-GCTGCCGATGAGCAGATGCTG
NAAT-49A3 Fwd-AACGTCAGCTCCTTTCCTGATGAT
NAAT-49A3 Rev-AGCAGCAGCCAGCATCTGCTCATCTT

The GST-importin α2 NAAT mutants 28A4, 39A5, and 49A were generated by inserting the *Bam*HI–*Eco*RI PCR fragments amplified from pEGFP plasmids into the *Bam*HI–*Eco*RI sites of the N-terminal GST-fused protein expression vector pGEX-6P-2 (GE Healthcare, US). Recombinant proteins were obtained as described in Supporting Methods 1. The plasmids for GST-importin α1 (18), GST-importin α2 (4), and GST-importin α3 (19) were obtained as previously described.

The fragments from the upstream regulatory region (LOC108961160 on chromosome 17, NCBI Reference Sequence: NG_051978.1) of mouse POU5F1 (MGI:101893, NCBI Gene: 18999) were obtained as follows. The target fragments were obtained from Bruce-4 cell genome DNA with the following targeting primers:

EcoRV upstream-1: Fwd-TACGATATCCACATCTGTTTCAAGCTAGTTCTA
EcoRV upstream-1-1: Rev-TACGATATCTGAATCTTCCGTTTCCTCC
EcoRV upstream-2: Fwd-TACGATATCGAGAATTATCAGGAGTTCAAGG
EcoRV upstream-2-1: Rev-TACGATATCACTTCCTGCTCCCCA

The obtained fragments were inserted into pBlueScript vector, followed by PCR using the primer sets

upstream-1 Fwd/upstream-1-2 Rev and upstream-2 Fwd/upstream-2-2 Rev,
upstream-1-2 Rev-TGAATCTTCCGTTTCCTCCA
upstream-2-2 Rev-ACTTCCTGCTCCCCAACC

### Genomic DNA shearing

Bruce-4 cells grown at 60%-70% confluence on 0.1% gelatine-coated 10-cm culture dishes were lysed in ChIP Elution Buffer (50 mM Tris-HCl pH 8.0, 10 mM EDTA, 1% SDS), and were sonicated six times for 5 s at an output level of 4 with Handy Sonic (TOMY, Japan), followed by seven cycles of 30 s on and 60 s off at the high level with Bioruptor II (BM Equipment, Japan). The cell lysates were then incubated for 3.75 h with 250 mM NaCl and 0.25 mg/mL Proteinase K (Nacalai Tesque Inc., Japan.). The sheared genomic DNA was purified using the FastGene Gel/PCR Extraction Kit (Nippon Genetics Co., Ltd., Japan) and fractionated by electrophoresis in 2% agarose gel. The fraction containing 600-bp genomic DNA was cut from the agarose gel (cutting images shown in Supporting Fig. S1) and underwent DNA purification using the FastGene Gel/PCR Extraction Kit (Nippon Genetics Co., Ltd., Japan).

### Gel-shift assay

The interaction of the importin α2 protein and DNA duplex was analysed by the gel-shift assay. The probes were 600-bp duplex DNA fragments (upstream-1, upstream-2, and genomic DNA) with biotinylated 3′ ends. DNA biotinylation at the 3′ end was performed using the Biotin 3’ End DNA Labeling Kit (Thermo Fisher Scientific, US) in accordance with the manufacturer’s instructions with some modifications. For this step, DNA duplex fragments at a final concentration of 0.236 pmol were biotinylated with terminal deoxynucleotidyl transferase (TdT) in 50 μL of a labelling reaction mixture (1X TdT Reaction Buffer, 0.5 μM Biotin-11-UTP, 0.15 U/μl TdT) at 37°C for 30 min, which was followed by termination by addition of 2.5 μL 0.2 M EDTA and TdT removal from the labelling reaction mixture by chloroform extraction. The biotinylated DNA fragments were applied to the gel-shift assay by using the LightShift Chemiluminescent EMSA Kit (Thermo Fisher Scientific, US) and Chemiluminescent Nucleic Acid Detection Module (Thermo Fisher Scientific, US) according to the manufacturer’s instructions. In this step, importin α2 recombinant proteins at a final concentration of 17.24 pmol or 34.48 pmol and 0.33 μL of the labelling reaction mixture containing biotinylated DNA duplex fragments were mixed to allow binding in a 20-μL binding reaction mixture (1X Binding Buffer, 50 ng/μL, Poly (dl·dC), 2.5% glycerol, 0.05% NP-40, 5 mM MgCl_2_). Then, the binding reaction mixture was subjected to electrophoresis at a 200-V constant pressure current in 4% TBE polyacrylamide gel. The DNA fragments in the gel were transferred to the nylon membrane and were probed with Streptavidin-Horseradish Peroxidase Conjugate. The Streptavidin-Horseradish Peroxidase Conjugate was detected using Pxi4 (Syngene, UK) or G:BOX mini (Syngene, UK). All assays were tested three times in independent experiments (See also Supporting Methods and Fig. S1).

### Chromatin immunoprecipitation

Chromatin immunoprecipitation assays were performed as previously described (20) with some modifications. 1.0 × 10^7^ ES cells were crosslinked in 0.5% formaldehyde containing media for 5 minutes at room temperature followed by termination of the reaction with 0.125 M glycin. Cell pellets were then suspended in RIPA buffer (50 mM Tris-HCl [pH 8.0], 150 mM NaCl, 2 mM EDTA, 1% NP-40, 0.5% sodium deoxychorate, and protease inhibitor cocktail) and were sheared for seven cycles of 30 s on and 60 s off at the high level with Bioruptor II (BM Equipment, Japan). The cell lysates were then diluted to a DNA concentration of 100 ng/μL after centrifugation at 15000 × *g* for 10 minutes and were applied to IP using specific antibodies. 25 μL of the diluted cell lysate for each sample was used as inputs in later steps. 1 μg of the antibodies, anti-KPNA2 (importin α2) (rabbit polyclonal, Abcam, UK) or normal rabbit IgG (EMD Millipore Corp) as a control were independently added to 250 μL of cell lysate solution with 20 μL of Dynabeads M-280 (Thermo Fisher Scientific, US). The IP solutions were incubated at 4°C overnight with rotation, and the beads were washed with ChIP buffer (10 mM Tris-HCl pH 8.0, 200 mM KCl, 1 mM CaCl_2_, 0.5% NP-40, and protease inhibitor cocktail), wash buffer (10 mM Tris-HCl pH 8.0, 500 mM KCl, 1 mM CaCl_2_, 0.5% NP-40, and protease inhibitor cocktail), and TE buffer. Reverse crosslinking was achieved by mixing the beads with 50 μL of ChIP Elution Buffer (50 mM Tris-HCl pH 8.0, 10 mM EDTA, 1% SDS), and the input lysate was applied similarly after adding 25 μL of ChIP Elution Buffer followed by incubation with 2% proteinase K (Nacalai Tesque Inc., Japan.) at 50°C for 1h. Obtained DNA was purified using the Wizard SV Gel and PCR Clean up system (Promega, US) in a final volume of 50 μL. See also Supporting Methods 2.

### Quantitative PCR

Quantitative PCR assays after ChIP were performed using THUNDERBIRD SYBR qPCR Mix (TOYOBO, Japan) and the Thermal Cycler Dice Real Time System II (TAKARA BIO, Japan) according to the manufacturer’s protocols with the primers below. In brief, two of the 50-μL eluted solutions obtained in ChIP were used for each reaction with 0.6 μL of 10 μM primer solution and 10 μL of THUNDERBIRD SYBR qPCR Mix with nuclease-free water (Nacalai Tesque Inc., Japan) up to 20 μL in total. Reaction cycles were as follows: 95°C for 30 s, 95°C for 15 s/54°C for 15 s/60°C for 30 s (cycled, Ct values under 37 were judged to be valid), 95°C for 15 s/60°C for 30s, 95°C for 15s (single cycle to obtain the dissociation curve). The standard curves for either primer sets were obtained using the genomic DNA purified from input lysates with four dilution rates, x0, x0.1, x0.01, x0.001, for individual samples. The experiments were triplicated, and the mean value of the importin α2 ChIP sample relative to one for control samples using normal rabbit IgG were calculated from the standard curve. All Ct values for the relative quantity in importin α2 ChIP samples were within the range of the standard curves obtained. Then, the relative ratio was calculated as (importin α2 ChIP)/(ChIP control). Four independent experiments were performed. The target sequences were upstream of the mouse gene POU5F1 (MGI: 101893, NCBI Gene: 18999, LOC108961160 on chromosome 17, NCBI Reference Sequence: NG_051978.1.)

upstream-1: Fwd-CACATCTGTTTCAAGCTAGTTCTAAGAA
upstream-1-2: Rev-CAACCTTGTCTTATGGATTGTTCTCTT
upstream-2: Fwd-ATGAAGACTACCATCAAGAGACACC
upstream-2-2: Rev-TTGTCTGTCTGCTCCTACACCAT

### Homology modelling of N-terminal domain of importin α2 protein

Homology modelling of the importin α2 IBB domain was performed using the Swiss Model with default parameters. Amino acid sequences of the IBB domain of mouse importin α2 were applied as target sequence, and PDB ID: 1QGK was selected as the template.

### Circular dichroism (CD) spectroscopy

Far-ultraviolet CD spectra from 300 nm to 195 nm were collected on a Jasco J-820 spectropolarimeter at room temperature with 50 mM importin α peptide in 20 mM phosphate buffer at pH 7.0 by using a 0.05-cm path-length fused silica cuvette. A band path of 1-nm and the corresponding scan speed were selected. Sixty-four scans were averaged, and the blank spectrum was subtracted from the sample spectra to calculate ellipticity.

### NMR measurement

NMR spectra were acquired on a Bruker Avance II 800 MHz spectrometer equipped with a cryogenically cooled proton optimised triple resonance NMR ‘inverse’ probe (TXI) (Bruker Biospin, Germany). All spectra were acquired at 25°C (298 K) with 256 scans using the p3919gp pulse program. Acquisition and spectrum processing were performed using Topspin3.2TM software. Chemically synthesised peptide and DNAs were dissolved in 50 mM Tris-Cl buffer solution containing 10% D_2_O, pH 7.9 and used for the measurements in 4-mm*ϕ* Shigemi tubes. The amino acid sequence of the chemically synthesised peptide was KDSTEMRRRRISNVELRKAKKDEQMLKRR (importin α 2_21-50), and the 15-bp DNA sequences were GCA GAT GCA TAA CCG (SOX-POU) and GCG GAC CAC TAG ACG (randomised control sequence).

### Treatment of transfected HeLa cells with Triton X100 and DNase I

HeLa cells were seeded on a coverslip and transfected with the GFP-fused constructs at an equimolar concentration by using Lipofectamine 2000 in accordance with the manufacturer’s protocol (Thermo Fisher Scientific, US). After 24 h, the cells were treated with Triton X100 and DNase I according to a previous report (17). In brief, cells were washed with buffer (20 mM HEPES-KOH [pH 7.3], 110 mM CH_3_COOK, 5 mM CH_3_COONa, 2 mM (CH_3_COO) 2Mg, 2 mM DTT, 1 μg/mL of leupeptin, pepstatin, and aprotinin), and treated with 0.5% Triton X-100 for 5 min at 4°C in the same buffer. The cells were washed with the buffer and were incubated with 0.2 mg/mL DNase I (Worthington Biochemical, US) for 1 h at 37°C. After fixation with 3.7% formaldehyde in PBS, cells were incubated with 1 μg/mL Hoechst (Nacalai Tesque Inc., Japan) for 5 min at room temperature. The images of GFP-fusion proteins were obtained by confocal microscopy A1 (Nikon, Japan) equipped with a 60X objective lens using the software NIS-Elements (Nikon, Japan). For quantification of nuclear GFP-fusion proteins, the phase-contrast microscope CKX53 (OLYMPUS, Japan) equipped with a 20X objective lens and software Cell Sens Standard was used under constant conditions for each experiment. Images obtained from two independent experiments were analyzed by Image J (NIH, US). The nuclei were identified as Hoechst-stained particles and plotted as ROIs (regions of interest), followed by quantification of nuclear distribution by measuring the mean value of fluorescence intensity of GFP image for each nucleus. The mean values of background intensity were subtracted for each sample. Tukey HSD Test were performed for statistical analysis. (https://astatsa.com/OneWay_Anova_with_TukeyHSD/).

## Results

### Importin α2 interacts with the genomic DNA of ES cells

Among importin α family proteins, importin α2 is highly and predominantly expressed in mouse ES cells and is involved in important physiological functions. To evaluate the chromatin-related functions of importin α2, we first assessed the nuclear distribution of importin α2 in mouse ES cells. Endogenous importin α2 was localised both in the cytoplasm and the nucleus (Fig. 1*A*), as shown by immunostaining assays using two different antibodies. Widespread nuclear staining of importin α2 was observed. A similar staining pattern has been reported in HeLa cells, in which the distribution of nuclear importin α2 diminished with DNase treatment (17). If the nuclear distribution could be attributed to direct interaction with DNA, the widespread staining indicates that importin α2 has the ability to directly bind to a relatively wide range of DNA sequences. Thus, we tested whether and to what extent importin α2 binds to genomic DNA to assess the possibility of a direct interaction with chromatin DNA. We prepared mixtures of genomic DNA fragments with sizes of approximately 600 bp by digestion of chromatin from undifferentiated ES cells and assayed whether importin α2 binds to these DNA mixtures by performing gel-shift assays (Figs. 1*B*, S1*A*-*C*). In the gel-shift assays, despite its low isoelectric point at 5.49, importin α caused apparent shifts of the DNA band in a dose-dependent manner. The results clearly indicated that importin α2 has affinity for diverse sequences of genomic DNA because not only parts of the mixture but the overall fractions of 600-bp fragments were also shifted by the addition of wild-type importin α2.

**Fig. 1.**
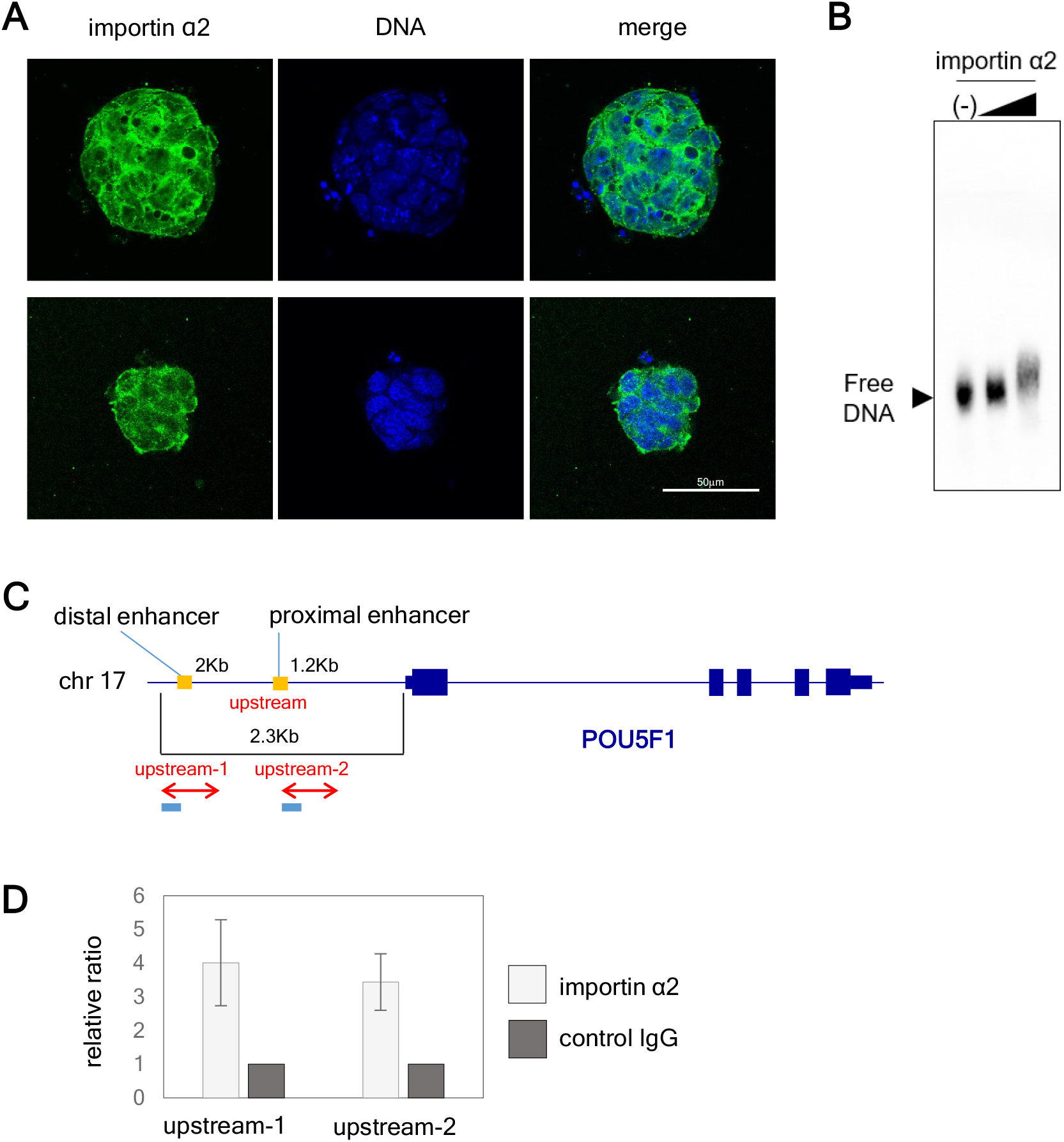
Importin α2 localises in the nucleus and binds the genomic DNA, including the upstream region of the POU5F1 gene, with affinity for diverse sequences in undifferentiated mouse ES cells. (A) Endogenous importin α2 localisation was shown by immunofluorescence in undifferentiated ES cells using two different antibodies. Upper panel: goat polyclonal antibody, lower panel: rat monoclonal antibody, green: importin α2, blue: DNA. (B) Sheared genomic DNA of approximately 600 bp was purified from undifferentiated mouse ES cells and used for gel electrophoresis with recombinant importin α2 protein. The black triangles above the gels indicates the two concentrations of importin α2 added to the binding reaction, 17.24 or 34.48 pmol as final concentrations. (C, D) ChIP-qPCR analysis with a specific antibody for importin α2 or the non-specific control IgG were performed to confirm the binding of importin α2 to the POU5F1 gene. (C) The primer sets used in the experiments are shown. (D) Results of ChIP-qPCR for the POF5F1 primers upstream-1 and upstream -2. The measured quantity of the importin α2 IP sample against a ChIP control sample is listed in terms of mean values of relative ratio with error bars (SE) in four independent experiments. Relative ratios were calculated and shown as importin α2 ChIP/ChIP control (shaded bar), ChIP control/ChIP control (=1, closed bar). See also Supporting Fig. S1.

Next, to determine whether importin α2 binds genomic DNA *in vivo*, we performed chromatin immunoprecipitation quantitative PCR (ChIP-qPCR) analyses using undifferentiated ES cells. Since the accessibility to DNA is affected by the structure of chromatin, which depends on the cell stages, we chose two target sequences whose activities are expected to be different in undifferentiated ES cells. We termed the two 200-bp DNA sequences of mouse POU5F1 (Oct3/4 gene) upstream regions as ‘upstream-1’ and ‘upstream-2’, where upstream-1 included the conserved distal enhancer CR4 domain and upstream-2 included the proximal enhancer domain. These two enhancers differentially contributed to stage-specific Oct3/4 expression during embryogenesis (21). The distal enhancer is known to be essential and sufficient for Oct3/4 expression in ES cells while the proximal enhancer is rather necessary for epiblasts (22). Primer sets to amplify the first 200 bp of each upstream region were used in importin α2 ChIP-qPCR (Fig. 1*C*). The results showed that both upstream-1 and upstream-2 were stably detected after positive PCR amplification from importin α2 ChIP samples (Figs. 1*D*, S1*D*-*G*). Taken together, these results suggested that multiple DNA sequences, including the POU5F1 genomic region, potentially interact with importin α2. Therefore, we further adopted these regions to identify the DNA binding domain of importin α2.

### Importin α2 directly binds the DNA via the IBB domain

To clarify the mechanism of importin α2 binding to DNA, we decided to determine the molecular binding mode *in vitro* by using mouse genomic DNA. First, we checked whether the importin α2 protein directly binds to DNA by performing gel-shift assays using the recombinant importin α2 protein and the double-stranded DNAs corresponding to the mouse ES cell genomic sequence (Figs. 2 and S2). The two upstream 600-bp DNA sequences of the POU5F1 gene tested in ChIP-qPCR assays (Fig. 1*C*) were used in this assay. The wild-type importin α2 interacted with both upstream DNA fragments of the POU5F1 gene, confirming that the DNA fragment from this region effectively bound importin α2 *in vitro*.

**Fig. 2.**
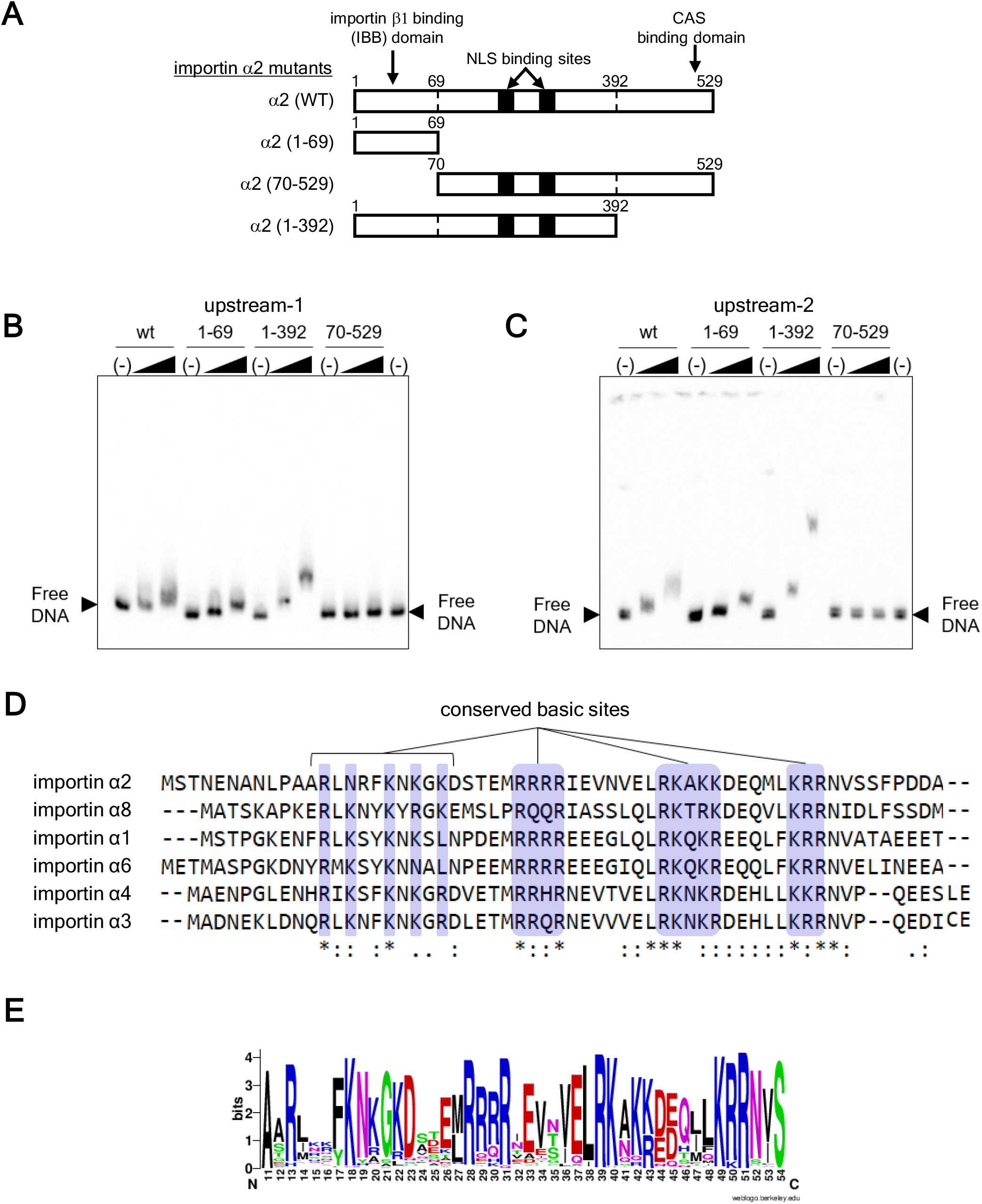
Importin α2 bound DNA *in vitro* via the IBB domain. (A) Full-length and deleted-recombinant importin α2 used in gel-shift (*in vitro* binding) assays. (B-C) The genomic DNA sequence from the importin α2-bound region in Oct3/4, determined by the ChIP-qPCR analysis in Fig. 1C and D, was selected and applied to these *in vitro* binding assays. DNA sequences from the POU5F1 upstream region (upstream-1 and -2) were located as described in Fig. 1C. Naked DNAs of upstream-1 (B) and upstream-2 (C) were used. The black triangles above the gels indicates the two concentrations of importin a2 added, 17.24 or 34.48 pmol as final concentrations. See also Supporting Fig. S2. (D) Conservation of basic amino acid sites of the importin α IBB domains. Amino acid sequences of mouse importin α (KPNA) families were aligned using ClustalW (DDBJ). The basic amino acid clusters without gaps and a cluster with gaps are presented in blue boxes. (E) A graphical representation of the amino acid conservation (Sequence logo) of IBB domains created with WebLogo (see Supporting Methods 7.1.2. for detailed procedure). The height of each letter within the stack indicates the relative frequency of each amino acid at that position.

Next, we determined the DNA-binding region of importin α2 by conducting additional gel-shift assays using recombinant importin α2 mutant proteins lacking functional domains. We focused on three major parts of importin α2: (1) the IBB domain with a linker, (α2 [1-69]); (2) the main body (α2 [70-392]); and (3) the C-terminal region (α2 [393-529]) (Figs. 2*A* and S2*A*). The recombinant mutant proteins containing the IBB domain showed significant shift from the original free DNA for both upstream-1 and upstream-2 (Figs. 2*B*, *C*, S2*B*-*E*). These indicate that the proteins have affinity to both 600-bp DNA sequences of the POU5F1 gene. In contrast, the mutant lacking the IBB domain (α2 [70-529]) did not show any significant shift. Here, the IBB domain that has been characterized as a coordinator of nuclear transport and autoinhibition was also necessary for DNA binding of importin α2 *in vitro*. The velocity of migration in electrophoresis under a spatially uniform electric field reflects both the hydrodynamic particle size and the net charge. Although the IBB domain of importin α itself is positively charged (the isoelectric point is 10.61), the main body of importin α, especially its C-terminal portion, bears a high negative charge, and the isoelectric point of full-length importin α is 5.49. Consequently, the reduction of migration (shift) upon binding of full-length importin α to DNA sequences is attenuated because the negative net charge of importin α partly offset the reduction of the migration. On the other hand, the deletion mutant α2 (1-329) lacking the highly negatively charged C-terminal portion has an isoelectric point of 8.80, and this mutant can be expected to show relatively large reduction of migration (large shift) due to its high positive net charge.

### Basic amino acid clusters within the IBB domain contribute to the binding of importin α2 to DNA

To elucidate the portion of the IBB domain responsible for DNA binding, we also examined whether any sequence homology to known nucleic acid-binding proteins was present in the IBB domain sequence of importin α. A protein BLAST search against the protein and nucleic acid structure database (Protein Data Bank) revealed that only the importin IBB domain families showed significant homology to the whole region of the IBB domain of importin α2 (data not shown). However, a BLAST search under search parameters adjusted for short input sequences detected many short motifs in the database. DNA-binding and RNA-binding proteins were then extracted from the hit sequences, and these analyses revealed that 24,830 of the 156,365 PDB entries shared short fragments that showed significant homology to importin α2_22-51 (KDSTEMRRRRISNVELRKAKKDEQMLKRR) (see Supporting Methods 3 for detailed procedures). Surprisingly, importin α2, which had never been classified as a nucleic acid-binding protein, showed homology with a large number of nucleic acid-binding proteins, even though the matched sequences were only on short length scales. These matching sequences included 675 entries for complexes of DNA and protein and 799 entries for complexes of RNA and protein. These entries accounted for approximately 13% of all the DNA-protein complexes (5071 entries) in the PDB and about 38% of all RNA-protein complexes (2104 entries), respectively. These hits included 27 DNA-protein complexes containing a tetra-R motif which is also found in the IBB domain (Supporting Table S1-1) and 36 RNA-protein complexes containing the motif (Supporting Table S1-2). Thus, astonishingly, the importin α IBB domain, a well-known canonical domain for nuclear transport regulation, was found to possess short basic motifs similar to those in nucleic acid-binding proteins. Furthermore, a clear diversity was found in the interaction pattern of the tetra-R motif with the nucleic acids, in the protein secondary structures of the region containing the motif, and in the target nucleic acid structures and sequences (Supporting Table S1). The manner in which the tetra-R motif directly comes into contact with DNA was roughly divided into three types of interactions: 1) as a part of the α-helix that binds deeply to the major grooves of DNA; 2) as a part of the extended strand that binds to the minor grooves; and 3) as a part of the α-helix riding on nucleic acid phosphate backbones. Considering these results together, the tetra-R motif appeared to be a useful element in interactions with nucleic acids. On the basis of these results and the known affinity of basic amino acids to DNA (23), we hypothesised that importin α2 might have interacted with DNA via the basic amino acid-rich portion within the IBB domain. Then, we aligned the importin α family IBB domain to check for conservation of the basic amino acids in the IBB domain (Figs. 2*D* and *E*). While some variations in amino acid sequences were seen, four clusters of basic amino acids, including the tetra-R motif, were essentially conserved among the importin α family proteins.

Thus, we focused on the three prominent basic amino acid clusters, 28RRRR, 39RKAKK, and 49RRR, of the IBB domain and designed amino acid substitution mutants to clarify the importin α2 DNA-binding domain. The R and K residues within each cluster were substituted to A, as shown in Fig. 3*A* (see also Fig. S3*A*), to create the mutants 28A4, 39A5, and 49A3, respectively. The R- and K-to-A substituted recombinant mutant proteins were used in gel-shift analyses to test the binding affinity against the same POU5F1 gene DNA sequences. The shift was significantly abolished by the mutations, with the effect of the 28A4 mutation being especially salient (Figs. 3*B*, *C*, S3*B*-*E*). Furthermore, we examined the binding ability of the mutant to genomic DNA mixtures (Figs. 3*D*, S3*F*-*G*), and the mutants (28A4, 39A5 and 49A3) showed only a weak or no interaction.

**Fig. 3.**
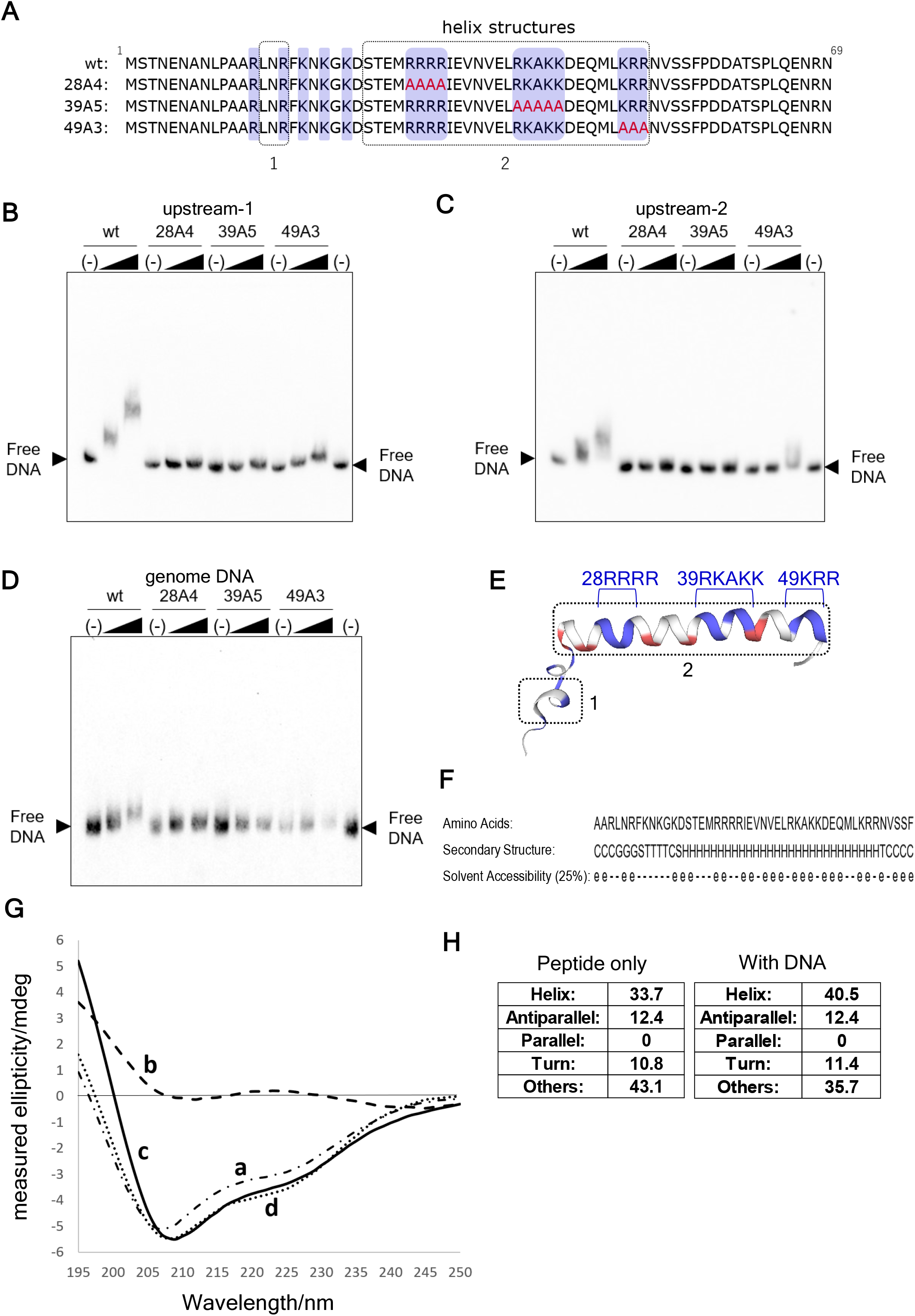
Importin α2 bound DNA via basic amino acids of the helical structure in the IBB domain. (A) The basic amino acid clusters in the IBB domain of importin α2 were mutated as indicated. The helical portion explained in (D) is indicated in box no. 1 and 2. (B-D) Recombinant wild-type and mutant importin α proteins were applied to *in vitro* binding assays using naked DNAs for upstream-1 (B) and upstream-2 (C) regions of the Oct 3/4 gene, and sheared genomic DNA of approximately 600 bp which was purified from undifferentiated mouse ES cells (D). The black triangles above the gels indicates the two concentrations of importin α2 added to the binding reaction, 17.24 or 34.48 pmol as final concentrations. (E) The location of the basic clusters indicated on the helix structures obtained by the Swiss Model with the template of IBB in the complex with importin β1 (PDB ID: 1QGK), where the basic amino acids are coloured blue and the acidic amino acids are coloured red. The box nos. 1 and 2 correspond to those presented in (A). (F) The amino acid sequence was analysed to predict the structural features through the web server SCRATCH. (G) CD spectroscopy to predict the helix content in IBB (mouse importin α2_ 1-69 peptide) in the absence and presence of dsDNA (upstream-1 600 bp). a) 6.6 μM of peptide only, b) 6.6 μM of DNA, c) mixture of the peptide and DNA, d) difference spectrum obtained by subtracting b) from c). (H) The predicted secondary structure contents for the peptide in the absence and presence of DNA. The left panel shows the structures in the absence of the DNA and the right panel shows the structures in the presence of the DNA. The unit of vertical axis is ellipticity described in mdeg. See also Supporting Fig. S3, 4.

### Importin α–DNA association model

We next attempted to determine how importin α2 binds to DNA. The amino acids 14L-16F and 24S-51R of the IBB domain adopt an α-helical conformation in the crystal structure of the complex with importin β1 (PDB ID: 1QGK) (Figs. 3*A* and *E*). In contrast to the crystal structures of the complex with importin β1, the IBB domain of importin α has been reported as a missing part in many crystal structures (for example, PDB ID: 5TBK; 5HUW; 5HUY; 5V5P; 5V5O; 5W4E). This suggests that the IBB domain adapts multi-conformational states or disordered conformation when it is not bound to importin β1. Therefore, we predicted the intrinsic propensity of the IBB domain for three-dimensional structure formation by analysing the amino acid sequence through the web server SCRATCH, which is based on machine learning methods (24). This analysis revealed that the central part 24S–51R of the IBB domain tends to easily form an α-helix structure, at least after being induced by other molecules (Fig. 3*F*). Accordingly, the results of CD measurements of the importin α2_1-69 peptide showed that the importin α2_1-69 peptide has an intrinsic helix-rich structure (estimated helix content, approximately 33.7%), and the addition of double-stranded DNA (upstream-1, 600 bp) slightly increased the helix content to 40.5% (Figs. 3*G*, *H* and S*4*). These values corresponded to that of the secondary structure predicted by SCRATCH, suggesting that the basic amino acid clusters within the IBB domain interact with DNA in an α-helical conformation as is in the crystal structure with importin β1 (9).

We next investigated the binding pattern of the IBB domain to DNA by docking and Molecular dynamics (MD) simulations using the α-helical structure of IBB in the complex with importin β1 (PDB ID: 1QGK). To predict the binding features of importin α2 and DNA, we constructed a model structure of the IBB domain α-helix-DNA complex using AutoDock vina (25) and analysed the α-helix-DNA interaction at the atomic level by performing a 30-ns MD simulation of the docked model structure to optimise the structures (see Supporting Methods 4 and Discussion for the computational experiment). The energetic and conformational analyses of the MD trajectory revealed two α-helix/DNA-binding modes (mode A and B) (Figs. 4*A*-*D* and S5). The α-helix in the IBB domain was fitted into a major groove of the canonical B-form DNA strands for mode A (Figs. 4*A* and *C*), while it was placed on DNA in parallel for mode B (Figs. 4*B* and *D*). In mode B, the peptide interacts with the phosphate backbone of the DNA at four points simultaneously, which is possible because the basic amino acids on the helix are concentrated on one side (Fig. S6 and Supporting Methods 5). These binding modes correspond to the two binding types for α-helical states out of the three tetra-R motif binding types mentioned in the BLAST search results in section 3. The electrostatic interactions between the negatively charged phosphate groups in DNA and the positively charged patches (28RRRR, 39RKAKK, and 49KRR) on the surface of the α-helix were important in both modes, and the α-helix did not directly read specific bases in the DNA. Interestingly, the α-helix moved on the DNA from state A to B during our short-term simulation.

**Fig. 4.**
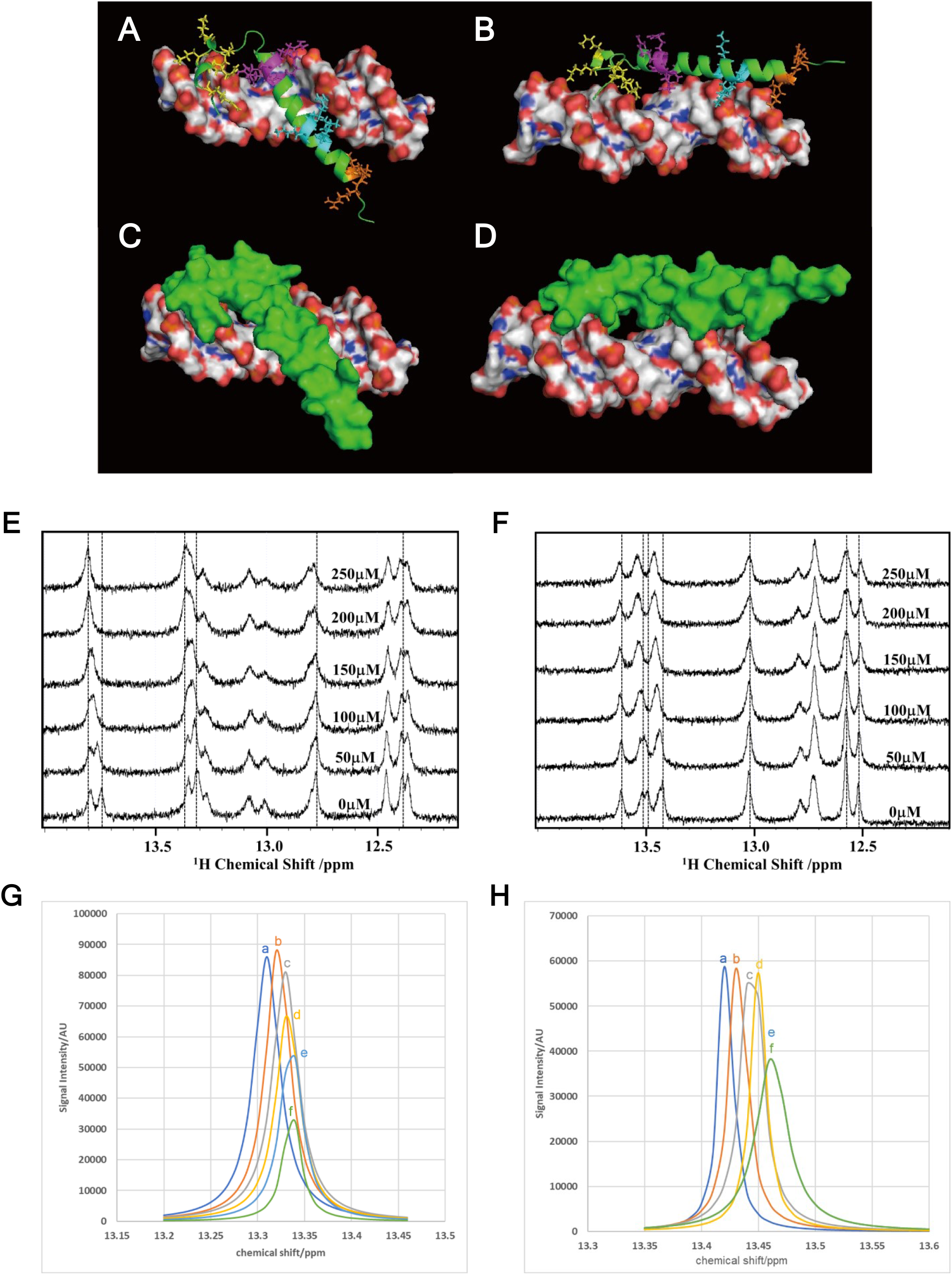
Importin α2 bound DNA in multi-mode. (A-D) Two model structures of the importin α2 IBB domain/DNA complex were obtained from molecular docking and MD (molecular dynamics) simulation: mode A (A and C) and mode B (B and D). DNA is shown in the surface model and coloured according to atoms (carbon: grey, oxygen: red, nitrogen: blue, phosphorus: yellow) (A-D). The α-helices of importin α are shown in a ribbon model (A and B) or a surface model (C and D) and are coloured green. Three positively charged patches in the α-helix, 28RRRR, 39RKAKK, and 49KRR, are shown in sticks coloured in magenta, cyan, and orange, respectively (A and B). N-terminal basic residues that formed short helixes (13R, 16R, 18K, 20K, and 22K) are also shown as sticks and coloured in yellow. (E, F) ^1^H-^1^D NMR titration experiments. The spectral changes in the imino proton region of the DNA induced by addition of indicated concentrations of peptide, importin α2_21-50, the core part of the arginine-rich region in mouse importin α2, are shown. The spectral change for 50 μM of the 15-bp double-stranded DNA of the SOX-POU binding region upstream of POU5F1(E). The change with 50 μM of a 15-bp double-stranded randomised sequence DNA(F). The peptide concentrations are indicated in each figure. (G, H) Spectral perturbations observed in the titration experiments. A representative perturbation profile for the NMR spectra of SOX-POU DNA by mouse importin α2_21-50 peptide (peak9) (G), and random-sequence DNA by mouse importin α2_21-50 peptide (peak9) (H). Peaks are reconstituted with parameters obtained from the decomposition. (G-a) 50 μM of SOX-POU core sequence DNA in the absence of the peptide, (G-b) DNA with 50 μM of mouse importin α2_21-50 peptide, (G-c) DNA with 100 μM of the peptide, (G-d) DNA with 150 μM of the peptide, (G-e) DNA with 200 μM of the peptide, (G-f) DNA with 250 μM of the peptide, (H-a) 50 μM of random-sequence DNA in the absence of the peptide, (H-b) DNA with 50 μM of the peptide, (H-c) DNA with 100 μM of the peptide, (H-d) DNA with 150 μM of the peptide, (H-e) DNA with 200 μM of the peptide, and (H-f) DNA with 250 μM of the peptide. See also Supporting Figs. S5, 7.

We assessed the validity of our model structures (modes A and B) by introducing mutations (mutations 28A4, 39A5, and 49A3) in the positive electrostatic patches on the α-helix surface for both A and B mode structures and repeating the experiment. The calculated binding free energies indicated weakening of the binding affinities for these variants, reflecting the disappearance of a part of the positive electrostatic patch on the α-helix surface (Table 1).

**Table 1.** Calculated binding free energies (Δ*G*_bind_) between IBB domain and DNA in the MD simulations.

| $\Delta G_{\text{bind}}$ (kJ/mol) | Mode A<sup>a</sup> | Mode B<sup>b</sup> |
| --- | --- | --- |
| Wild type | -526.05 ± 36.28 | -602.96 ± 27.91 |
| 28A4 | -296.06 ± 40.04 | -431.71 ± 20.42 |
| 39A5 | -342.54 ± 10.21 | -426.85 ± 15.31 |
| 49A3 | -432.00 ± 21.46 | -508.82 ± 19.56 |

<sup>a</sup> Values for wild type was averaged over a 7 ns period ranging from 11 to 18 ns, those for variants were averaged over last 2 ns (8 ~ 10 ns).
<sup>b</sup> Values for wild type was averaged over a 5 ns period ranging from 25 to 30 ns, those for variants were averaged over last 2 ns (8 ~ 10 ns).

Furthermore, to elucidate the binding strength and stoichiometry of importin α to DNA, we performed one-dimensional NMR titration experiments of chemically synthesised 15-bp double-stranded DNA (GCA GAT GCA TAA CCG), which is the core sequence of the SOX-POU binding region in the upstream-1 region, including the POU5F1 distal enhancer, with chemically synthesised peptides (importin α2_21-50, the core part of the basic amino acid-rich region in mouse importin α2) (Figs. 4*E*, *F*, S7 and Table S2). The chemical shift changes in the imino proton region with the addition of peptide are shown in Fig. 4*E*. We also examined the interaction between importin α2_21-50 and a randomised sequence DNA duplex (GCG GAC CAC TAG ACG), which has the same length as the SOX-POU core sequence (Fig. 4*F*). In the NMR titration experiments, several peaks in the DNA imino proton region of 1D spectra were apparently perturbed by the addition of the peptide in both experiments (see Supporting Methods 6 for the detail). The observed spectral changes obviously showed two phases in each titration experiment, indicating that the binding of the peptide to DNA had at least two modes (Figs. 4*E*-*H* and S7, Table S2). Therefore, the spectral changes of the typical peaks in the entire imino proton region were analysed by non-linear least-square fitting to obtain the dissociation constants and stoichiometries for the binding by assuming two-step multivalent binding.

The estimated binding strengths were K_d1_ = 3 × 10^−7^ and K_d2_ = 6 × 10^−4^ with respect to the SOX-POU sequence, and the stoichiometries were 1:2 (DNA:peptide) for the first strong binding mode and 1:3 for the relatively weak binding mode. For the randomised sequence DNA duplex, the estimated apparent binding strengths were K_d1_ = K_d2_ = 1 × 10^−4^, and the change was saturated at a stoichiometry of 1:4 (DNA:peptide). The changes in the chemical shifts without remarkable broadening of resonance line widths (Figs. 4*E*-*H*) indicate that binding interactions occur in the fast exchange regime on the NMR chemical shift timescale, which in turn suggests that the binding dissociation constant K_d_ would have to fall in the 10^−6^ M range or more (weaker) (26). This held true for both modes of binding to randomised sequences, suggesting that the binding was almost diffusion-limited, whereas one of the estimated dissociation constants for SOX-POU was out of this range. This implies that the on-rate for binding to SOX-POU was significantly facilitated and that each microscopic step of the binding event had a short half-life, despite the relatively strong overall binding. Since multivalent binding with a relatively weak binding strength on each site resulted in an apparent intermediate-strength binding, the binding features can be listed as follows: the binding is multivalent; both on-rate and off-rate are high; the apparent overall binding affinity is intermediate; and the binding is semi-specific for the DNA sequence (with some extent of sequence preference).

As described in Figs. 3*A* and 3*E*, the importin α IBB domain contains conserved basic amino acid sites around the N-terminal short helix (14L-16R in importin α2), 28RRRR, 39RKAKK, and 49RRR, and the sites touched DNA either in modes A or B in the MD simulation (Figs. 4*A*-*D*). Here, we designated the DNA-binding domain of importin α, with the four conserved basic sites in the IBB domain (13R-51R of mouse importin α2), as the “Nucleic Acid Associating Trolley pole” (NAAT) domain. Note that the NAAT domain does not include the NLS-binding sites nor is it located close to these sites; therefore, importin α should be capable of binding DNA and cargo simultaneously.

### Importin α bound DNA and was retained in the nucleus via the NAAT domain in cells

We next attempted to examine the binding of the NAAT domain to DNA in cells. We first tested the effect of the NAAT 28A4 mutation, which clearly extinguished DNA-binding ability in the gel shift assay (Figs. 3*B*-*D* and S3*B*-*G*), on the localisation of GFP-fused importin α2 protein. The wild-type GFP-importin α2 localised in the nucleus and partially in the cytoplasm (Figs. 5*A*, S8). On the other hand, GFP-28A4 was mainly observed in cytoplasm with weak staining in the nucleus (Figs. 5*A* and S8), suggesting the role of NAAT for nuclear retention. Since importin α is a nucleocytoplasmic shuttling protein, and since the NAAT domain overlaps with IBB domain, alterations in the efficiency of nuclear entry or exit and nuclear retention may be coupled with nuclear localisation of the 28A4 mutant. Thus, we decided to conduct experiments using export-deficient importin α2 mutants.

**Fig. 5.**
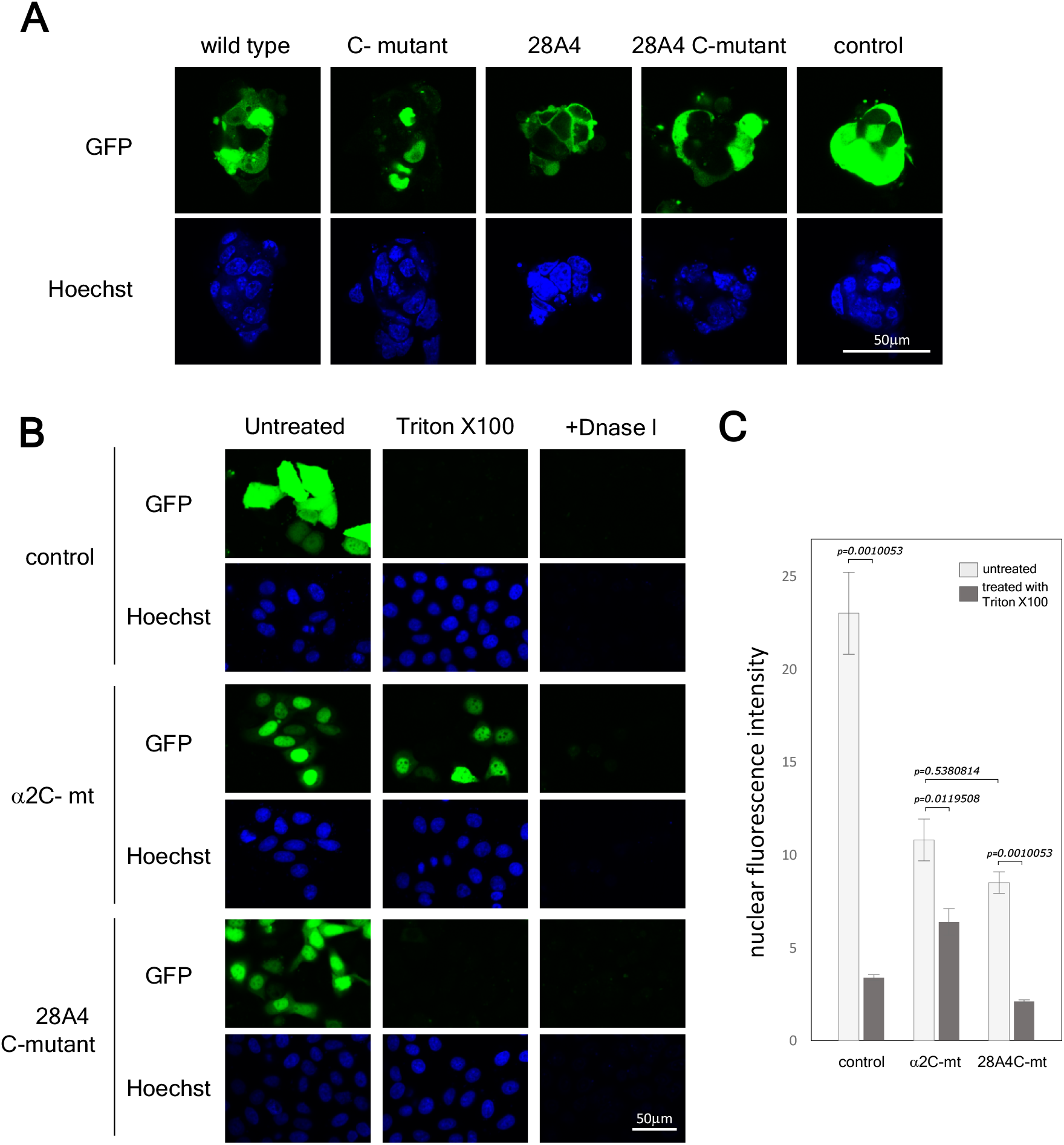
Importin α2 bound DNA via the NAAT domain in cells. (A) GFP-fused importin α2 proteins were expressed in ES cells. GFP-wild-type importin α2, 28A4 mutant, importin α2 C-mutant, and 28A4 C-mutant are shown with control GFP (green), as indicated. The blue colour of Hoechst staining indicates DNA. (B) HeLa cells transfected with GFP-importin α2 C-mutant, GFP-28A4 C-mutant, or control GFP (green) were treated with Triton X100 or Triton X100 followed by DNase I treatment. See also Fig. S8. (C) The nuclear localisation of the GFP-importin α2 C-mutant and GFP-28A4 C-mutant with or without Triton X100 treatment were measured. Fluorescence intensity in the nuclear area was plotted. The numbers of samples are as follows: control (n = 142), control/Triton X100 (n = 113), GFP-importin α2 C-mutant (n = 221), GFP-importin α2 C-mutant /Triton X100 (n = 210), GFP-28A4 C-mutant (n = 189) and GFP-28A4 C-mutant/Triton X100 (n = 152). The level of statistical significance is shown as the p value.

A mutation in the CAS-binding site in the C terminus, which has been established to accumulate importin α in the nucleus as a result of disruption of the nuclear export of importin α (17), was introduced to GFP-28A4. The result showed that the C-mutant enhanced nuclear localisation of both the wild-type and the 28A4 mutant of GFP-importin α2 (Figs. 5*A* and *B*). Next, to confirm the NAAT-DNA binding in cells, we treated HeLa cells with Triton X100 and DNase I and analysed the distribution of the GFP-28A4 C-mutant. Expectedly, a certain proportion of the nuclear GFP-importin α2 C-mutant was resistant to Triton X100 but was excluded by DNase I treatment (Figs. 5*B*), which was consistent with previously reported observations showing that nuclear localisation of endogenous importin α was resistant to Triton X100 but not to DNase I treatment (15, 17). On the other hand, although the GFP-28A4 C-mutant also abundantly localised to the HeLa cell nucleus, Triton X100 clearly diminished the nuclear fluorescence of this mutant to a comparable level to the control cells expressing diffusible GFP protein (Figs. 5*B*), suggesting again that NAAT binds DNA and retains GFP-importin α2 in cells. The nuclear signal of the GFP-28A4 C-mutant was slightly weaker but at comparable levels to the wild-type GFP-importin α2 C-mutant, and Triton X100 diminished the nuclear GFP intensity of the 28A4 C-mutant to around 25%, while the wild type retained 60% of its intensity (Figs. 5*C*). Although, in theory, nuclear migration could have been suppressed through a defect in importin β1 binding by the 28A4 mutation (27), this mutant has been shown to be capable of nuclear entry since the GFP-28A4 C-mutant accumulated in the nucleus before Triton X100 treatment (Figs. 5A, B and C). Note that importin α is known to migrate into the nucleus by itself without importin β1 (28).

On the basis of these results, we concluded that importin α2 binds genomic DNA via the NAAT domain not only *in vitro* but also *in vivo,* and that the NAAT domain promotes nuclear retention of importin α2 in cells. The heterogeneity of intracellular distribution of importin α proteins has been estimated to be a part of the regulatory mechanisms for its physiological functions, since nuclear localisation of family members is observed in specific stages of cellular events such as spermatogenesis (29), stem cell differentiation (30), and stress response (14–17). The direct binding of importin α to genomic DNA via the NAAT domain may play dominant role in this regulation.

### DNA-binding features appear to be conserved among importin α family proteins

The helix positions of the basic amino acids within the NAAT domain are fundamentally conserved among the importin α families (Figs. 2*D*, *E* and S*6*). Considering the fact that the basic amino acid clusters responsible for DNA binding of importin α2 are conserved, the DNA-binding feature via the NAAT domain seemed to be conserved among importin α family proteins.

In addition to basic amino acid conservation, one additional acidic amino acid is found in importin α3, 4, and two acidic residues were found in importin α1,6 with respect to α2,8 in the positively charged side of the helix. Thus, the importin α families can be characterized by the relative arrangement of the conserved basic amino acids, and these acidic amino acids were divided into three groups (Fig. S6). Unexpectedly, this grouping is consistent with the traditional classification of importin αs based on whole-length sequence similarity and their functions (11, 13). The coincident classification indicates that this grouping may reflect a relationship between the evolution of the canonical and non-canonical functions of importin α via the NAAT revealed in this study.

Such considerations prompted us to investigate whether the DNA-binding property is commonly conserved in the importin α family, and we used other importin α family proteins in the gel-shift assay with the same genomic DNA fragments as those used in the assays for importin α2. Actually, we used importin α1 and α3 as representatives of the subtypes. Both importin α1 and importin α3 shifted the whole fraction of the 600-bp sheared genomic DNA with minor differences in effectiveness (Figs. 6 and S9). Importin α3 bound with similar efficiency as importin α2, while importin α1 showed slightly reduced efficiency. These results clearly showed that importin α family proteins share DNA-binding characteristics. The results of the amino acid sequence analysis and the gel-shift assay indicate that the DNA-binding property is commonly conserved among importin α family proteins. However, the findings also suggest that there may be some differences in the DNA-binding property within the family as well. Detailed analyses of the differences are an interesting topic for future studies from the view of integrated functions of importin α family proteins for chromatin.

**Fig. 6.**
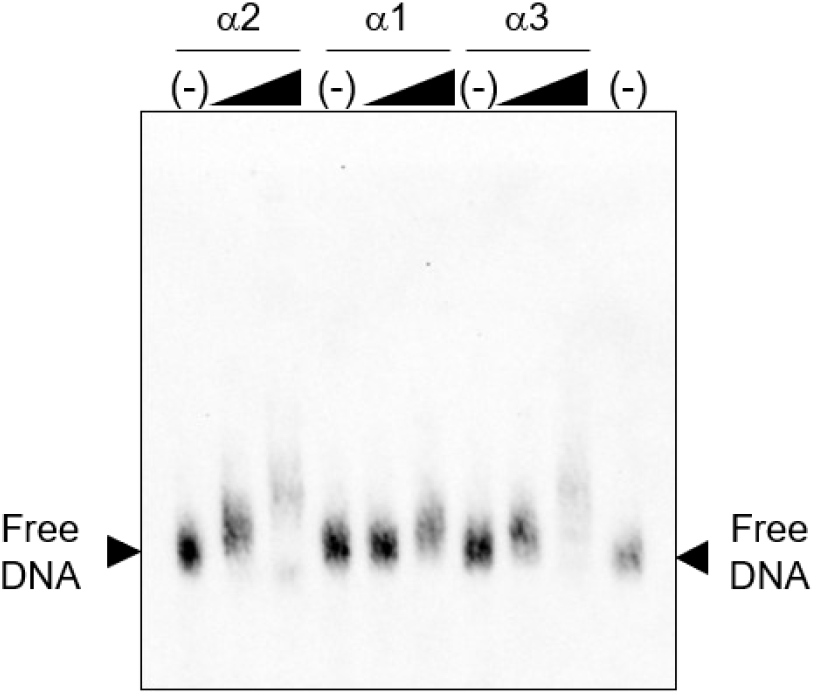
Importin α proteins multiply bound genomic DNA of ES cells. Sheared genomic DNA of approximately 600 bp purified from undifferentiated mouse ES cells was applied to gel shift assays with recombinant proteins of importin α2, α1, and α3, as indicated. The black triangles above the gels indicates the two concentrations of importin α2 added to the binding reaction, 17.24 or 34.48 pmol as final concentrations. See also Supporting Fig. S9.

## Discussion

The findings of this study suggest that importin α proteins can be revisited as DNA-associating proteins with unique characteristics. A novel DNA-binding NAAT domain was identified as a series of helix structures in the N-terminal IBB domain of importin α2 (Fig. 7*A*). The basic amino acids arranged along the helix as positively charged patches on the surface enable multi-mode binding of importin α2 to a broad range of genomic DNA molecules in a semi-specific manner, resulting in nuclear retention of importin α2, and this property is probably conserved among all importin α families.

**Fig. 7.**
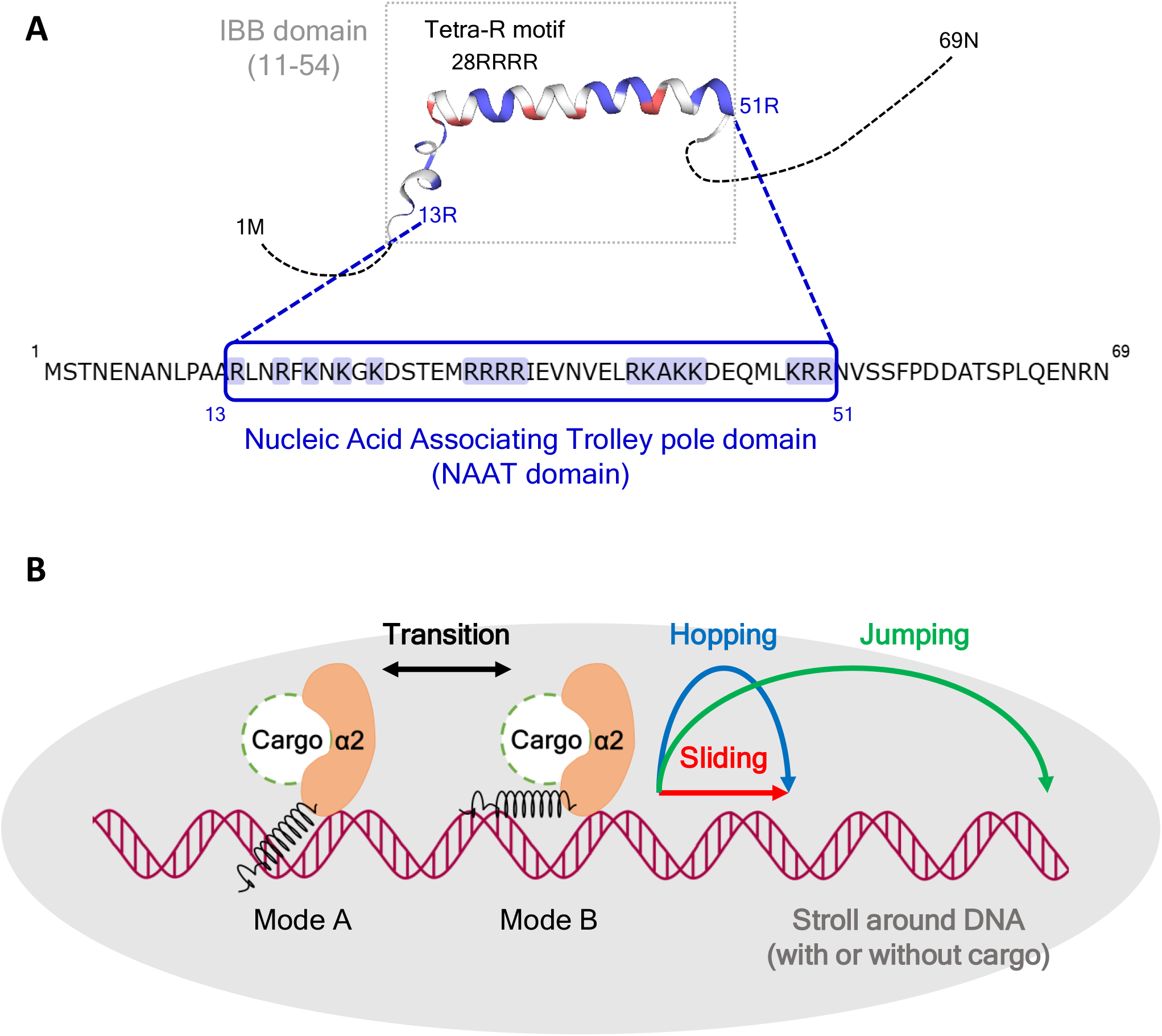
The importin α–DNA association model. (A) The schematic chart shows the NAAT domain of importin α2. The NAAT domain (13R-51R, blue box) localises within the IBB domain (dotted box) and contains basic amino acid clusters (shaded in blue), including the tetra-R motif (28RRRR) that predominantly binds DNA, which are indicated on the helix structures obtained by the Swiss Model with template of IBB in a complex with importin β1 (PDB ID: 1QGK) (11A-54S) adding the missing parts at both ends (1M-12A, and 52N-69N, dotted line), where the basic amino acids are coloured blue and the acidic amino acids are coloured red. (B) When importin α2 approaches chromatin (with or without cargo or interacting proteins), the Nucleic Acid Associating Trolley pole domain (NAAT domain, R13-K51 of mouse importin α2) in the helix structure within the IBB domain associates with DNA predominantly with a tetra-R motif. Importin α2 associates with DNA and strolls around by transitions among multiple modes, showing sliding, hopping, and jumping to neighbouring DNA. See also Supporting Fig. S11 for regulation of cargo fate determination.

The DNA-binding NAAT domain is distinct from the conventional DNA-binding domains of known DNA-binding proteins, although there are some common traits between NAAT and these proteins. For example, although the existence of positively charged amino acid clusters is a typical characteristic of DNA-binding proteins, as shown in our BLAST search, the semi-sequence–specific binding mode based on electrostatic interactions using basic amino acid clusters and the estimated tertiary structure of the NAAT complex are different from the known DNA-binding proteins. Most of the known sequence-specific protein-DNA associations are based on a system-specific fine balancing act of diverse competing forces and generally involve negligible to highly unfavourable net electrostatics, a highly favourable van der Waals interaction, and hydrophobic interactions, where the unfavourable net electrostatics for binding arise from the acidic and neutral residues despite the favourable contribution of the basic residues on the protein (31). Moreover, although NLS is mapped on the DNA-binding domains of many DNA-binding cargo proteins (32) and IBB/NAAT can also bind the NLS-binding site of importin α itself in an autoinhibitory manner (33), the NLS is generally placed out of the DNA-binding interface in the cargo proteins. In other words, the basic amino acid residues in the NLS of the cargos are independent of DNA-binding residues, while NAAT can bind to both DNA and the NLS-binding site using exactly the same residues. All these characteristics must be recognised as distinct structural traits in different lineages.

Our results show two distinct properties of the association between importin α and DNA (chromatin): first, importin α can interact with the vast majority of genomic sequences, and second, it simultaneously *shows* moderate sequence selectivity with preference for some specific sites. Consequently, the characteristics of the DNA-binding ability of importin α via the NAAT domain satisfy the necessary conditions for efficient delivery of itself and/or cargos to various sites of chromatin based on the framework of a dimension-reduced facilitated diffusion model for searching promoters of DNA-binding proteins on chromatin (34, 35). This physicochemical model describes that a certain DNA-binding protein randomly binds to multiple sites on the DNA and diffuses by sliding, jumping, or hopping along the DNA chain until it reaches its specific functional sites. This reduction in the dimensionality (3D to 1D) of the search space for the target enhances the search efficiency under certain conditions, and, in fact, some DNA-binding proteins have been known to be able to locate their target sites amid myriad off-target sequences within millions to billions of base pairs at remarkably rapid rates (36) that are sometimes two orders of magnitude larger than the value estimated from three-dimensional diffusion (37). Various theoretical and experimental studies have been conducted in order to characterize the mechanisms underlying this rapid target site search (38–48). Facilitated diffusion is one of the predominant models used to characterize this phenomenon (38). Although the concept of facilitated diffusion is a topic of debate (43, 46), the model has an increasing body of supporting evidence (46).

This mechanism necessarily requires the DNA-binding proteins to show semi-specific binding properties that are fundamentally non-selective but with some sequence preference for the target site (49,50), and the balance of these behaviours seems to determine the efficiency of the target search (46). It is also considered to be preferable that a protein (or a protein complex) has at least two DNA-binding surfaces or more to perform an intersegmental transfer from a DNA site to another site (43). These properties were exactly shown in the exchange of binding modes demonstrated or implied by MD simulation, biochemical mutant works, and NMR measurements for NAAT domain binding to DNA. Theoretical considerations also suggest that electrostatic interaction may play an important role in this facilitated diffusion (36, 43). For the facilitated diffusion of nearby off-target DNA sequences, a large positively charged patch on a protein surface is essential (36), and the sliding rate is influenced by the degree of positive charge clustering in the specific binding region of the nucleic acid-binding protein (51). Thus, the properties of the NAAT domain meet the requirements for facilitated diffusion, considering the four positively charged patches covering over a half of the entire surface (Fig. S6), and the fact that the results of the NMR titration experiments suggested that the process consists of relatively accelerated association and rapid dissociation as is described in section 4. In combination with the fact that the NAAT domain interacted with a wide range of sequences shown in the present study, the association of importin α with chromatin is suggested to follow the principle of facilitated diffusion. Regarding the binding specificity in the facilitated diffusion process, there is a trade-off between the search step efficiency and the recognition step, which usually exhibits relatively strong and specific binding (36). The results of NMR titration experiments apparently showed a sequence-dependent preference for the binding in addition to the non-sequence–specific binding ability. In addition, while transferring cargos including other DNA-binding proteins, the sequence specificity of importin α binding to DNA could be augmented by the cargos. Thus, these findings and the supporting evidence indicate that the binding properties of NAAT to DNA can facilitate both searching and recognising the target on the huge sequence space on chromatin, either directly or indirectly through facilitated diffusion, semi-specific association, and/or cooperative target recognition with cargos. Here, we propose the ‘stroll and locate’ model for the association of chromatin with importin α, in which importin α is continuously moving around DNA to rapidly search over a wide range of genome regions and arrive at the destination (Fig. 7*B*).

The multifunctionality of the IBB domain may also be important in this context. The non-canonical DNA-binding NAAT domain was identified in the canonical IBB domain required for nuclear transport of NLS proteins in an overlapping manner. For the canonical IBB domain, reports have described many intracellular functions of importin α by virtue of its binding to various proteins. For example, the interactions of the IBB domain with various proteins such as Nup50 (8), RAN (52), and RBBP4 (53) are known to be involved in the nuclear transport complex disassembly process, and the interaction with CAS (6, 54) is known to be involved in the export step since the IBB domain plays an important role for stabilization of the importin α/Cse1p (CAS)/RanGTP ternary complex (55,56). Furthermore, studies have investigated the relationship between the transport activities and formation of the autoinhibitory form in which IBB binds to the NLS-binding site of importin α itself, where the basic amino acids in IBB play a critical role (8–10). The functional amino acids involved in the interactions with the IBB/NAAT binding partners, including its autoinhibitory conformation, are summarized in Fig. S10 (also see Supporting Method 7). Since these binding sites partly overlap or line up closely, this overlapping may cause interference between the binding partners to some extent. However, importin α is likely to avoid such interference in vivo because of the distinct localisation of each binding partner in minute fractions in the nucleus. Furthermore, the NAAT-DNA interaction is predominantly electrostatic and seems to be different from the protein-protein interactions between IBB and other transport factors such as CAS and importin β. For example, IBB-importin β recognition occurs through a spatial arrangement of highly conserved acidic and hydrophobic residues (57), so the priority of the binding partner will be sensitively affected by the local microenvironment within the cell nucleus, such as the surrounding ion content, ion concentrations, and crowders, because changes in these conditions differently affect each type of interaction.

The destination of each component of the transport complex was shown to be stochastic after assembly and disassembly at nuclear pores, and the segregation was not strict in the cell nucleus (55). Even if the segregation of the binding partners is insufficient, the overlapping of the binding sites on the IBB will enable an enhanced sharp switching by reinforcement of the recombination of the complex, reflecting the differences in the concentration of the binding partners in each microcompartment and the differences in microenvironment as discussed above, and the final spatial regulatory mechanism may also be linked to structural regulation of chromatin.

The switching may be directly related to the successful cycling of importin α translocation from the cytoplasm to the nucleus, from the nucleoplasm to the DNA, and back from the nucleus to the cytoplasm. This switching feature may also explain previously reported data in which the nuclear transport function of importin α was altered by enhancement of nuclear accumulation-associated DNase-sensitive contents under stress conditions such as heat shock or oxidative stress (16), and which was used to suggest that dysfunction of importin α as a transport receptor was the dominant determinant of NLS transport suppression (58). Furthermore, the previous studies proposed that a mutual control mechanism exists in which the normal protein transport pathway is suppressed for the urgent transport pathways that facilitate the specific import of heat shock proteins (16,17) such as Hsp70 via Hikeshi (59). Thus, the overlap of canonical and non-canonical binding sites in the same domain, NAAT/IBB, seems to be a molecular basis for the functional coupling of suppression of nuclear transport and the nuclear retention in the stress response of cells.

Notably, the relationship between importin α chromatin association and transcriptional regulation is an important aspect to further understand the role of importin α-chromatin interaction. Importin α is known to carry various cargos, (13) including Oct3/4 (30,60–62), which binds to the cis element SOX-POU. The present study used ChIP-qPCR assays to demonstrate that importin α actually bound to the upstream sequence of the POU5F1 gene, and the NMR titration experiment showed that importin α bound to the SOX-POU element with greater binding preference than that for random-sequence DNA. These results imply that importin α not only carried transcriptional regulators in a non-sequence–specific manner, but also interacted directly with the target DNA sequences in a semi-specific manner probably to support the cargo protein binding. Interestingly, nuclear retention of importin α has been reported to coordinate HeLa cell fate through changes in STK35 gene expression (17). Moreover, a transcription factor Zac1 was shown to require importin α not only for the nuclear import but also for expression of the target gene p21 (63). In both studies, ChIP-qPCR assays found that the promoter regions of the targets, STK35 or p21, coprecipitated with importin α. Additionally, importin α forms complexes with the yeast transcriptional activator GAL4 and the GAL4 target DNA, independent of GAL4 nuclear transport (64). At least for these factors, it is worth investigating whether importin α is involved in their control via NAAT. From the perspective of cargo, the fates will be regulated by binding competitions and/or by cooperation among several factors, including IBB/NAAT, in the process of nuclear transport (see Fig. S11 for the cargo fate determination steps in detail). For example, TRIM28 is reported to possess importin α–dependent NLS which overlaps with the HP1 binding box, suggesting that the preferential interaction of TRIM28 to HP1 associated with heterochromatin occurs after the importin α–dependent nuclear transport (65). In combination with our results, this process may occur in a cooperative manner with NAAT to DNA binding and TRIM28 to HP1 binding.

In studies using *dim-3* (defective in methylation -3) mutants of *Neurospora crassa* in which importin α showed an E396K mutation, importin α was reported to be essential for heterochromatin formation and DNA methylation by targeting cargo proteins to chromatin (66,67). Interestingly, *dim-3* mutation decreased the interactions between constitutive heterochromatic domains. Since this amino acid is required for IBB autoinhibitory binding to importin α itself, the E396K mutation in *Neurospora* may influence the behaviour of IBB/NAAT and alter the interaction of importin α and chromatin. Another intriguing topic is the tethering of importin α to the membrane by palmitoylation during mitosis, which controls the balance of the importin α/β complex and thereby scales the mitotic spindle (68). The expanded perspective suggests that NAAT will play a role in the mitotic chromatin behaviour by sequestering importin α.

Among a large number of importin α–dependent cargos, there are hundreds of cargos that are related to transcription, chromatin organization, RNA processing, and nucleic acid metabolic processes (69). In the homology search for the IBB domain in this study, we identified many proteins in RNA-protein complexes possessing similar basic amino acid cluster as in NAAT (Supporting Table S1-2). Thus, the NAAT/IBB domain may bind not only to DNA but to RNA as well. Considering this information with the other findings, the binding of the importin α NAAT domain to chromatin may partly affect the structural changes of chromatin coupled with cargo proteins such as chromatin remodelling factors. Therefore, clarification of the relationship between NAAT and this nuclear sub-compartmentalization via phase separation is an important topic for future research.

The findings obtained in this study suggest that importin α plays pivotal roles in a wide range of delicate regulatory processes with subtlety in each step of nuclear events as a coordinator for delivering chromatin-binding proteins to their targets (Fig. S11). Although each step of this sequence should be reckoned with in detail in future, the use of NAAT peptides itself as a competitor should be a great advantage in further studies of physiological significance and drug discovery, considering the situation that more than 60 peptide drugs have already been approved for therapeutic use, and several hundreds of novel therapeutic peptides are under preclinical and clinical development (70).

Since all members of the importin α family seem to possess an NAAT domain and have slightly different amino acid sequences, variations in the physiological effects of importin α chromatin association via NAAT domain are also of considerable interest. The expression patterns of importin α family members show marked differences in different cell types, which indicates their roles in cell fate determination. For example, regulation of development and alterations in the expression profiles of different importin α subtypes during differentiation in ES cells and during spermatogenesis (reviewed in 29,71-72), etc. have been studied so far. Furthermore, different importin α proteins have different cargo specificities. Therefore, the interwoven variations in these factors could enable rigorous regulations. Conversely, the interactions between importin α and chromatin may also be modified by the presence of cargo proteins such as transcriptional regulatory factors, and these modifications may be responsible for the regulation of cellular physiological functions. If the chromatin association is influenced by its cargos, the composite effects of the variation of importin α types and of the cargos may be related to general transcriptional regulation and the variety of physiological functions.

In summary, we propose the ‘stroll and locate’ model to explain the association between importin α and chromatin via the NAAT domain after entering the nucleus (Fig. 7*B*), and we suggest that this model may be related to a wide variety of cellular physiological processes.

## Supporting information

supplementary discussions

## Funding

This work was supported by the Japan Society for the Promotion of Science (JSPS) KAKENHI to Noriko Yasuhara (18H04870, 15K07069, 25116008) and Noriko Saitoh (18H05531, 18K19310) and by a Nihon University Multidisciplinary Research Grant for 2018.

## Acknowledgments

We thank Drs. Naoki Horikoshi, Saki Hirata, Yuichiro Semba, S.J. Nogami, Hiroyuki Taguchi, Hitoshi Kurumizaka, Yasuyuki Ohkawa and Hiroshi Kimura for kind suggestions and discussions about this work. The constract for GST-importin α2 mutant (1-329) were kindly gifted from Dr. Yoshinari Yasuda.

## Notes

### Competing Interest Statement

The authors have declared no competing interest.

### Summary of Updates

Figures added.

