## supplementary figures for "Importin α2 association with chromatin: Direct DNA binding via a novel DNA binding domain"

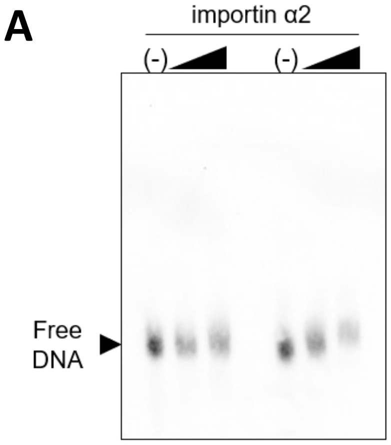


**Figure S1.** (A) The same experiments as in Figure 1B were performed independently. The black triangles above the gels indicates the two concentrations of importin α2 added to the binding reaction, 17.24 or 34.48 pmol as final concentrations. (B, C) Sheared Bruce-4 genomic DNA was applied to electrophoresis in a 2% agarose gel. Then, 600-bp DNA fractions were cut from the gel as indicated by the yellow boxes which the two of four samples are indicated as examples. (D, E) Samples of importin α2 ChIP were applied for quantitative PCR (q-PCR) using specific primers for the POU5F1 gene upstream region as described in Figure 1C. Four independent experiments (No. 1~4) were performed (D), and the mean ratio of sample/control were calculated (E), which is presented graphically in Figure 1D. (F) DNA electrophoresis was performed to check DNA fragmentation of the lysate used in ChIP analysis. (G) ChIP samples before protein digestion were applied to SDS-PAGE followed by western blot analysis to detect importin α2. Loading volume was 1/125 of the lysate used for IP. Arrowhead is the expected position of the importin α2 band.

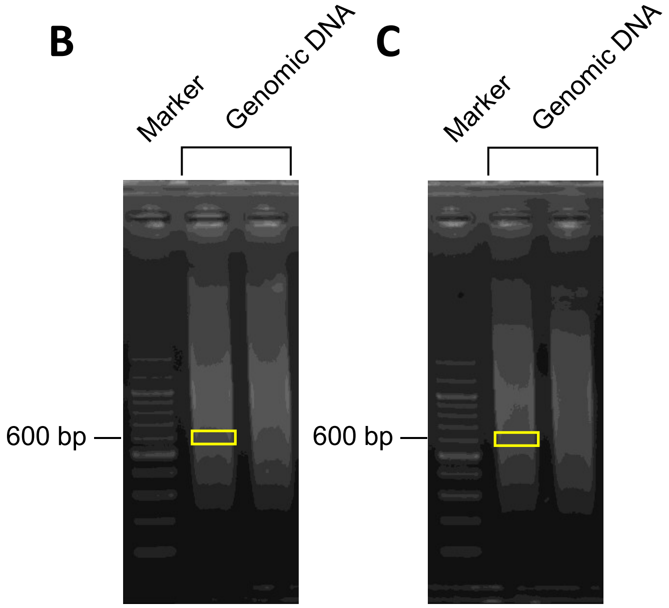


D

| experiment no. | target | sample | Rel. Qty |
|---|---|---|---|
| 1 | upstream-1 | normal IgG | 7.39E-03 |
| | | importin α2 | 1.20E-02 |
| | upstream-2 | normal IgG | 5.17E-03 |
| | | importin α2 | 8.64E-03 |
| 2 | upstream-1 | normal IgG | 7.71E-03 |
| | | importin α2 | 6.10E-02 |
| | upstream-2 | normal IgG | 8.64E-03 |
| | | importin α2 | 4.31E-02 |
| 3 | upstream-1 | normal IgG | 6.70E-01 |
| | | importin α2 | 1.22E+00 |
| | upstream-2 | normal IgG | 1.66E-01 |
| | | importin α2 | 3.07E-01 |
| 4 | upstream-1 | normal IgG | 4.19E-10 |
| | | importin α2 | 1.96E-09 |
| | upstream-2 | normal IgG | 1.92E-09 |
| | | importin α2 | 1.01E-08 |

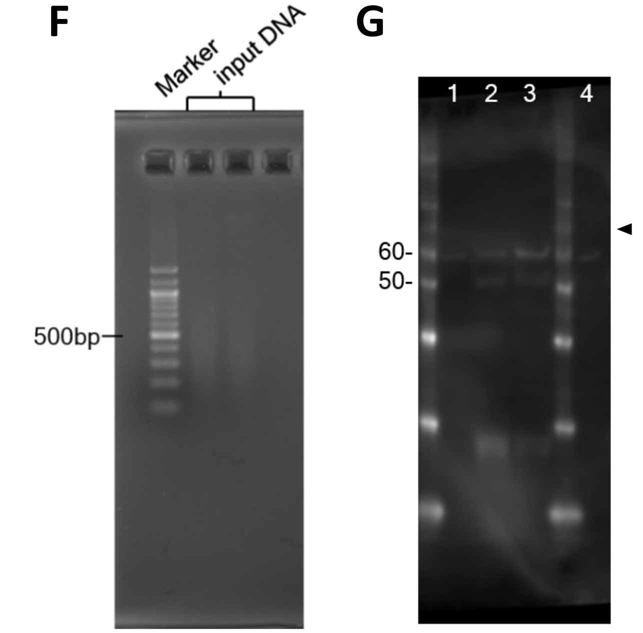


E

| ratio | upstream-1 | upstream-2 |
|---|---|---|
| sample mean | 4.010503781 | 3.43658916 |
| control | 1 | 1 |
| sample SE | 1.279026256 | 0.84007415 |
| control SE | 0 | 0 |

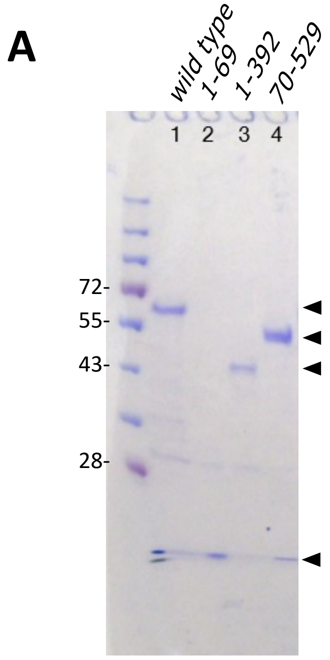


**Figure S2.** (A)Recombinant proteins of wild-type and deletion mutants of importin α2 used in this study were applied to SDSPAGE followed by CBB staining. (B-E) The same experiments as in Figure 2B, C were performed twice independently. The genomic DNA sequence from the importin importin α2 bound region in POU5F1 gene, determined by the ChIP-qPCR analysis in Figure 1B and C, were selected and applied to these *in vitro* binding assays. DNA sequences from the POU5F1 gene upstream region (upstream-1 and -2) were located as described in Figure 1B. Naked DNAs of upstream-1 (B, C) and upstream-2 (D, E) were used. The black triangles above the gels indicates the two concentrations of importin importin α2 added to the binding reaction, 17.24 or 34.48 pmol as final concentrations.

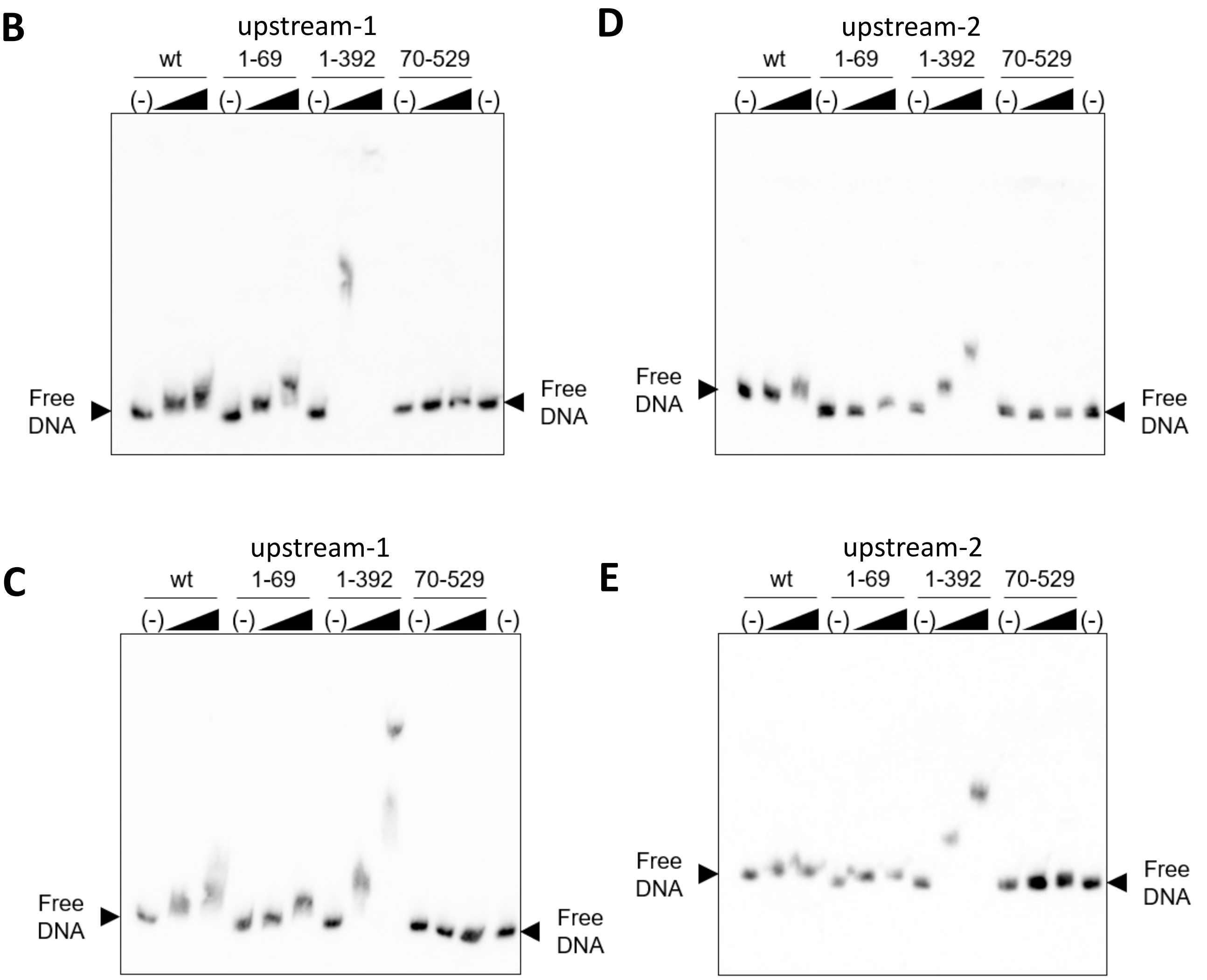

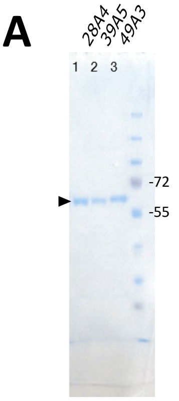


**Figure S3.** (A)Recombinant proteins of importin α2 NAAT mutants as indicated were applied to SDSPAGE followed by CBB staining. (B-E) The same experiments as in Figure 3B, C, H were performed twice independently. The genomic DNA sequence from the importin α2-bound region in POU5F1 gene, determined by the ChIP-qPCR analysis in Figure 1B and C, were selected and applied to these *in vitro* binding assays. DNA sequences from the POU5F1 gene upstream region (upstream-1 and -2) were located as described in Figure 1B. Naked DNAs of upstream-1 (B, C) and upstream-2 (D, E) were used. Importin α2 NAAT domain bound multiple genomic DNA sequences of ES cells. Sheared genomic DNA of approximately 600 bp purified from undifferentiated mouse ES cells was applied to gel electrophoresis with recombinant proteins as indicated (F, G). The black triangles above the gels indicates the two concentrations of importin α2 added to the binding reaction, 17.24 or 34.48 pmol as final concentrations.

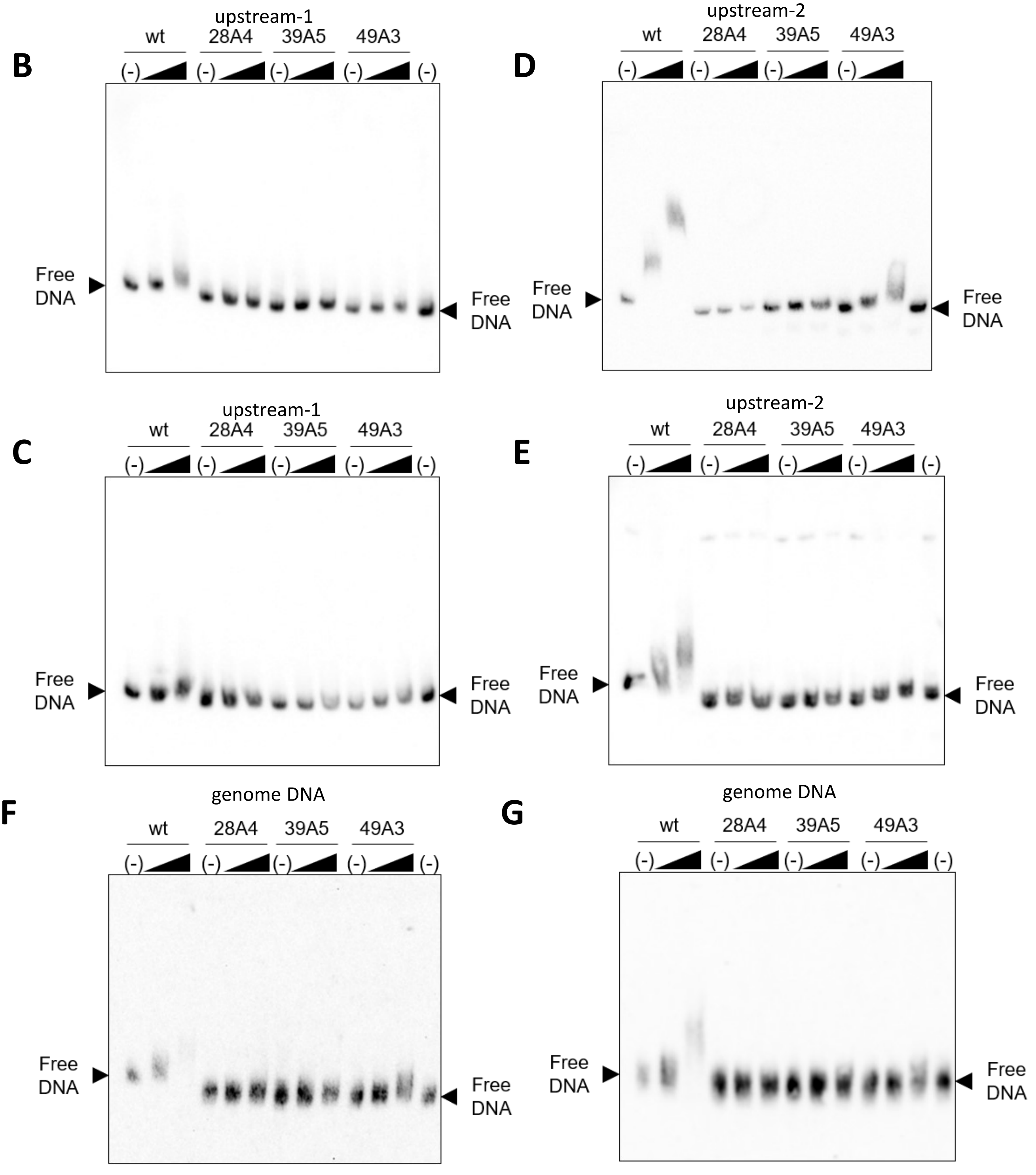

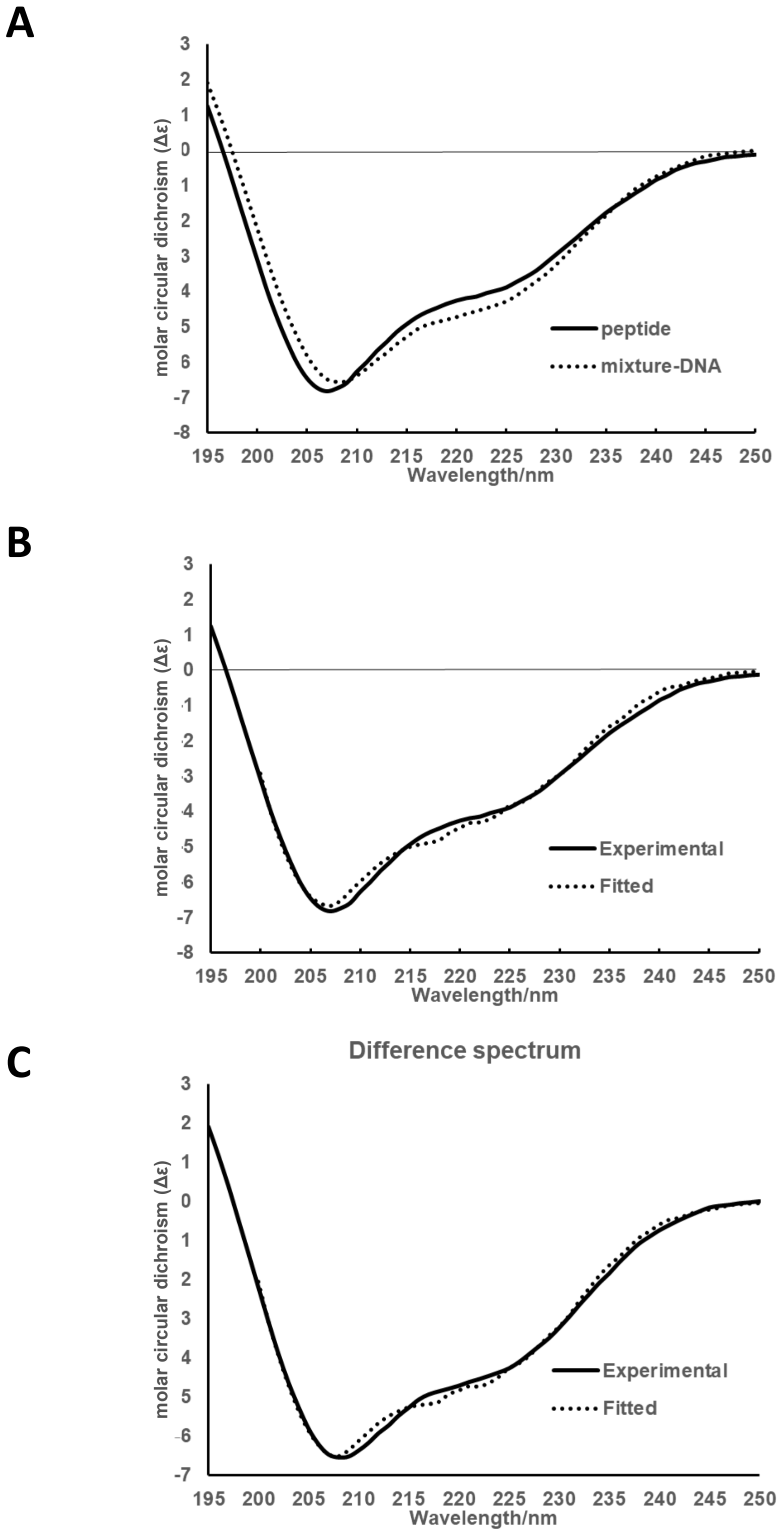


**Figure S4.** (A-C) Effect of DNA binding on the CD spectrum of importin α2_1-69 peptide (IBB domain of mouse importin α2). (A) CD spectra of importin α2_1-69 peptide in the presence of DNA (indicated as mixture-DNA) and in the absence of DNA (indicated as peptide). The spectrum of mixture-DNA was obtained by subtraction of the DNA spectrum b in the main text from spectrum c, shown in Figure 3F. (B) CD spectrum of importin α2_1-69 peptide in the absence of DNA (indicated as Experimental) and the fitted spectrum by BESTSEL (1) (indicated by Fitted). (C) CD spectrum of importin α2_1-69 peptide in the presence of DNA (indicated as Experimental) and the fitted spectrum by BESTSEL (indicated by Fitted). The unit of the vertical axis is molar circular dichroism (Δε) calculated using the mean residue concentration (CMR).

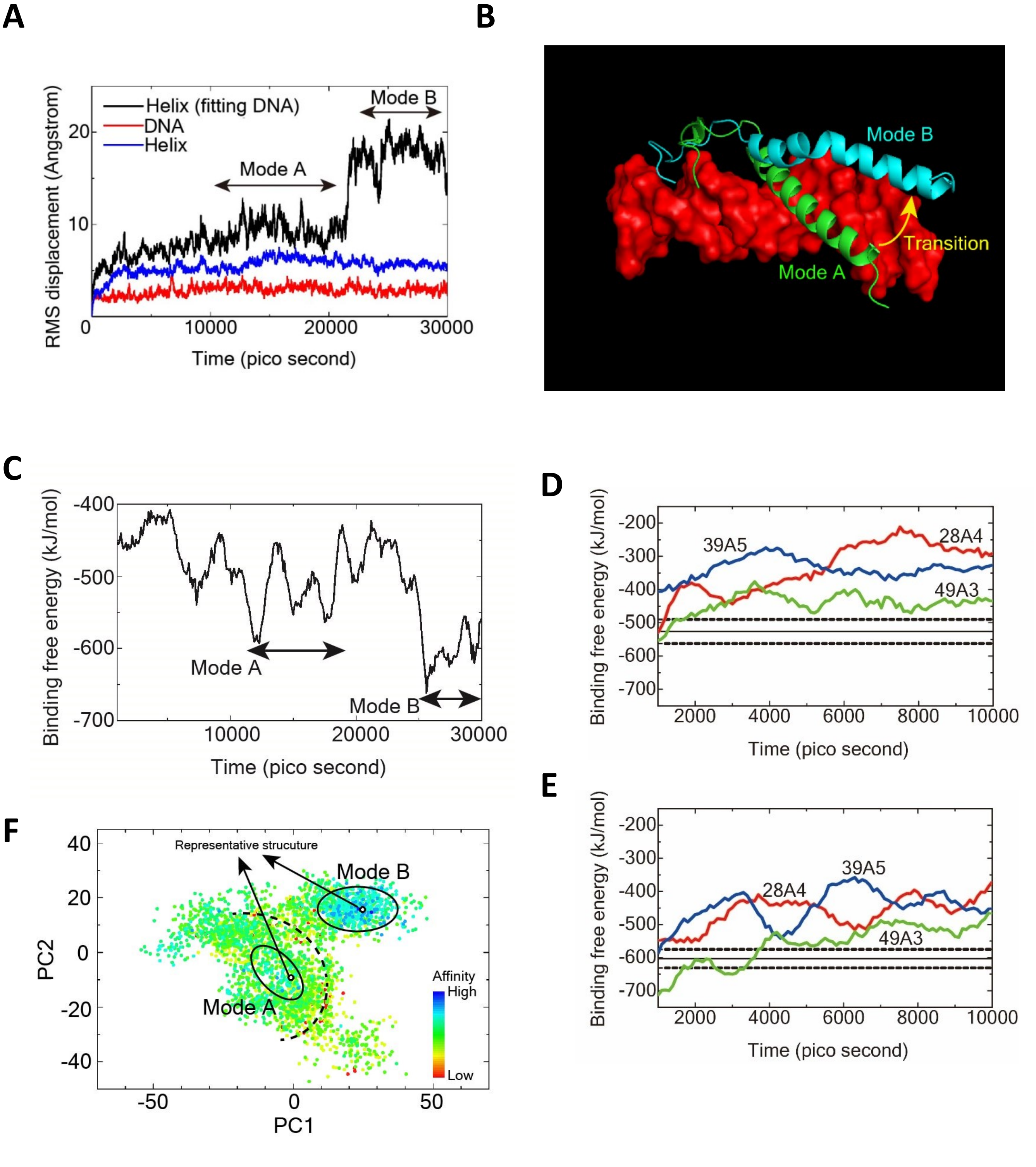


**Figure S5.** (A) Time-dependent RMS deviations of backbone heavy atoms from the initial docked model structure. (B) Comparison of binding modes of A (green) and B (blue). Double strand DNA is shown in red. (C) Time course of α-helix/DNA-binding free energy. (D, E) Time courses of α-helix-mutant/DNA-binding free energies for mode A (D) and B (E). The solid black lines show the averaged binding free energy values of the wild type, and the dashed black lines show the standard deviations. (F) Principal component analysis of the FEL for the α-helix/DNA complex. Binding free energies are mapped as a function of PC1 and PC2. Dashed black line shows the energy barriers in between binding modes A and B.

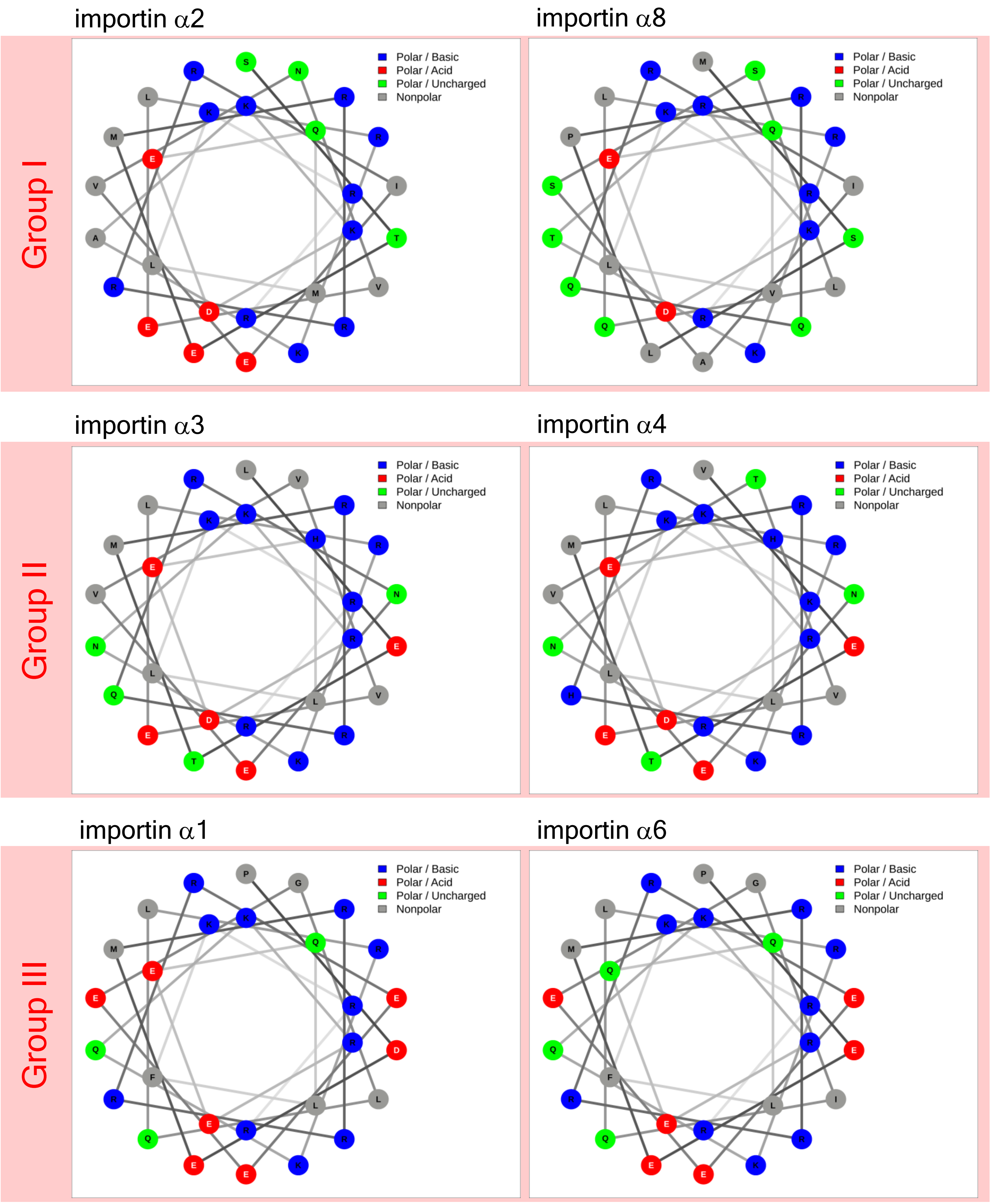


**Figure S6.** Basic amino acids forming positively charged patches are mostly located on one half (right side) of the surface of the α-helix in the NAAT domain corresponding to importin α2 (24S-51R). The importin α family is divided into three groups, group I with no acidic amino acid in the basic (right) side, group II with one, and group III with two.

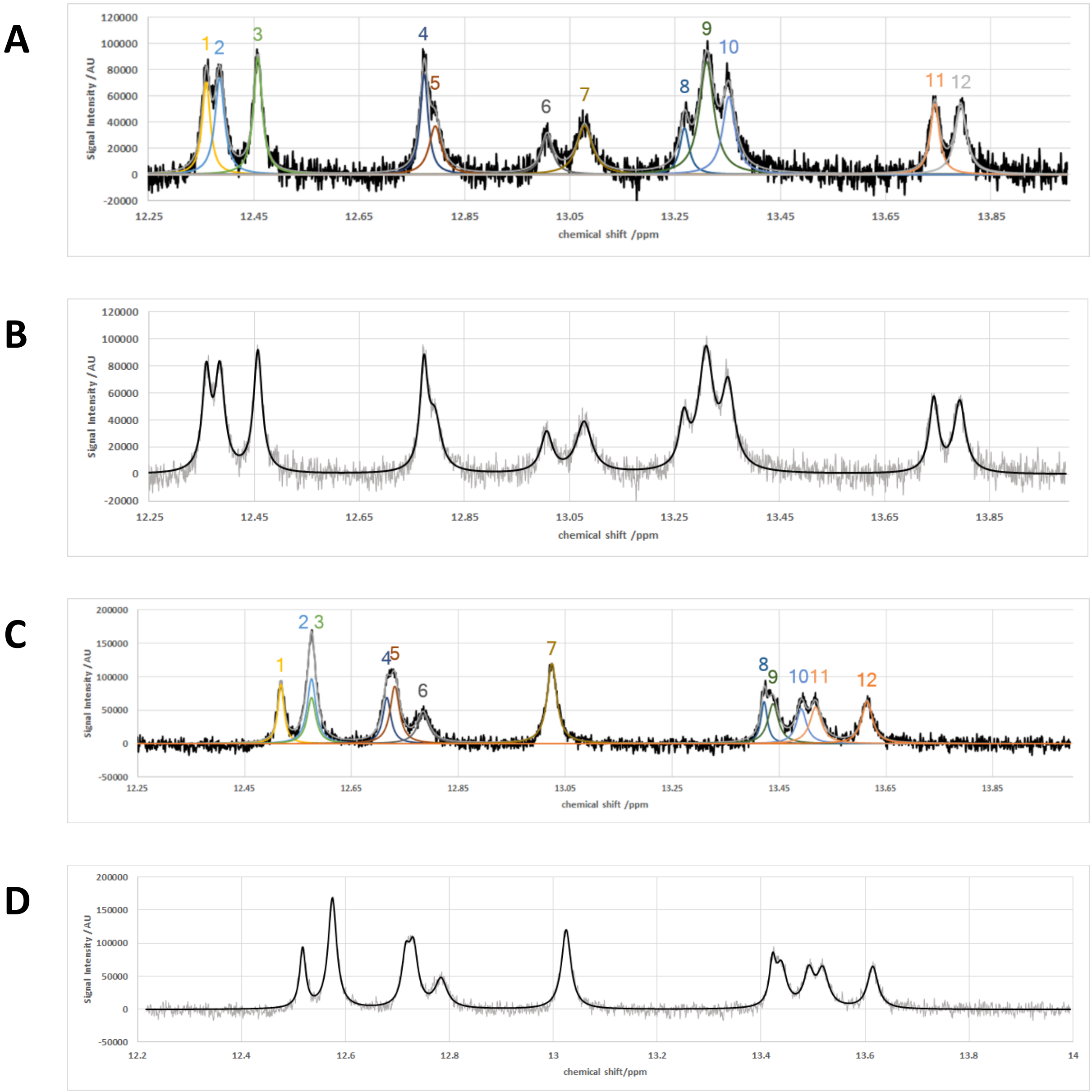


**Figure S7.** (A-D) Decomposition of the imino proton region of the NMR spectrum of SOX-POU core sequence DNA. Imino proton region of the NMR spectra for SOX-POU core sequence 15-bp duplex DNA (A, B) or for random-sequence 15-bp duplex DNA (C, D) were decomposed into twelve peaks by non-linear fitting using Lorentz peak functions. The spectrum in the absence of the peptide is shown representatively. (A,C) The observed spectrum (black) and the 12 decomposed components. (B, D) The observed spectrum (grey) and the spectrum reconstituted from the twelve components (black).

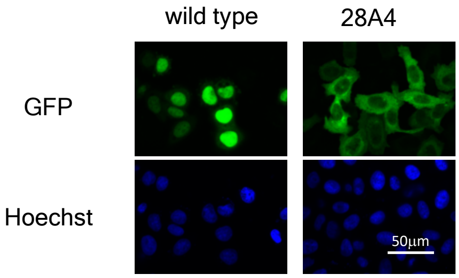


**Figure S8.** GFP-fused importin $\alpha$ 2 proteins were expressed in HeLa cells. GFP-wild type importin $\alpha$2, 28A4 mutant are shown with control GFP (green) as indicated. Blue colour of Hoechst staining shows DNA.

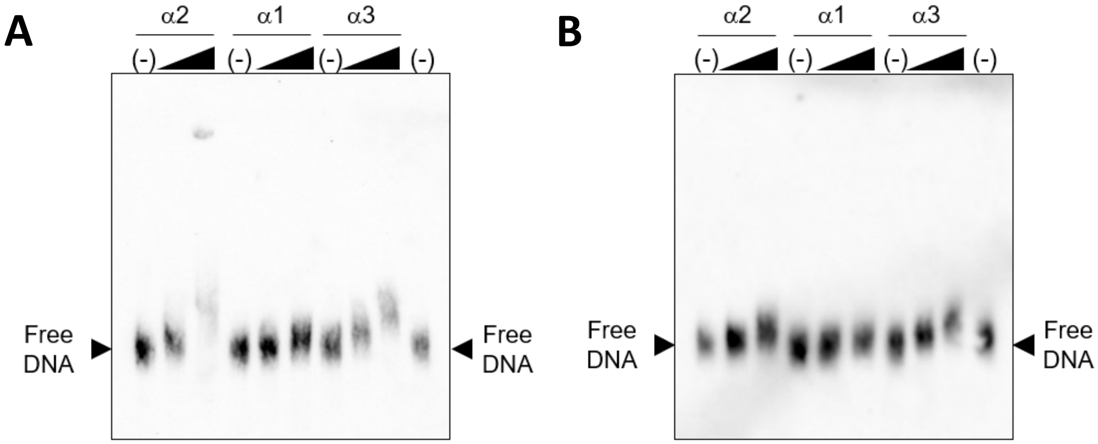


**Figure S9.** (A-B) The same experiment as in Figure 5 was performed twice independently. Importin α family proteins multiply bound genomic DNA of ES cells. Sheared genomic DNA of approximately 600bp purified from undifferentiated mouse ES cells was applied to gel electrophoresis with recombinant proteins as indicated. The black triangles above the gels indicates the two concentrations of importin α2 added to the binding reaction, 17.24 or 34.48 pmol as final concentrations.

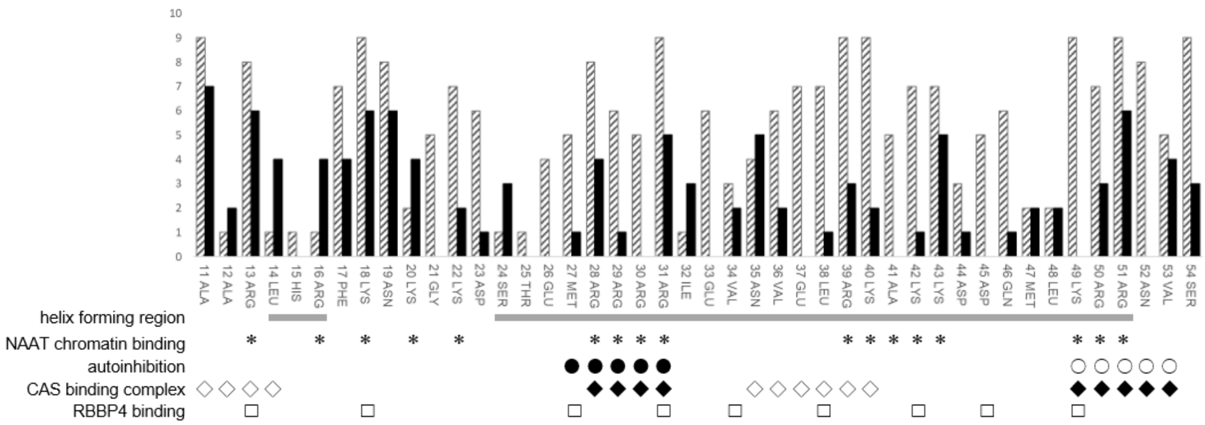


**Figure S10.** Molecular phylogenetic analyses of the IBB domain. The striped bar shows the conservation index and the black bar shows the relative accessible surface area change upon complex formation with importin β1. Functional amino acids in the IBB domain are marked below. The helix formed in the crystal structure of importin β1 bound to the IBB domain of importin α (PDB:1QGK) is underlined. The basic amino acids in the NAAT domain are marked by asterisks. The amino acids that bind to the major NLS-binding site in the crystal structure of mouse importin α2 (PDB: 1IAL) are marked by open circles. The amino acids that bind to the minor NLS-binding site in the crystal structure of rice importin α1α (PDB: 4B8J) are marked by closed circles. The amino acids that bind to either of the major and minor NLS-binding sites or that bind to CAS (Cse1p in yeast) in the crystal structure of the yeast exportin Cse1p complexed with importin α and RanGTP (PDB: 1WA5) are marked by closed diamonds or by open diamonds, respectively. The putative RBBP4-binding amino acids (1) are marked by squares.

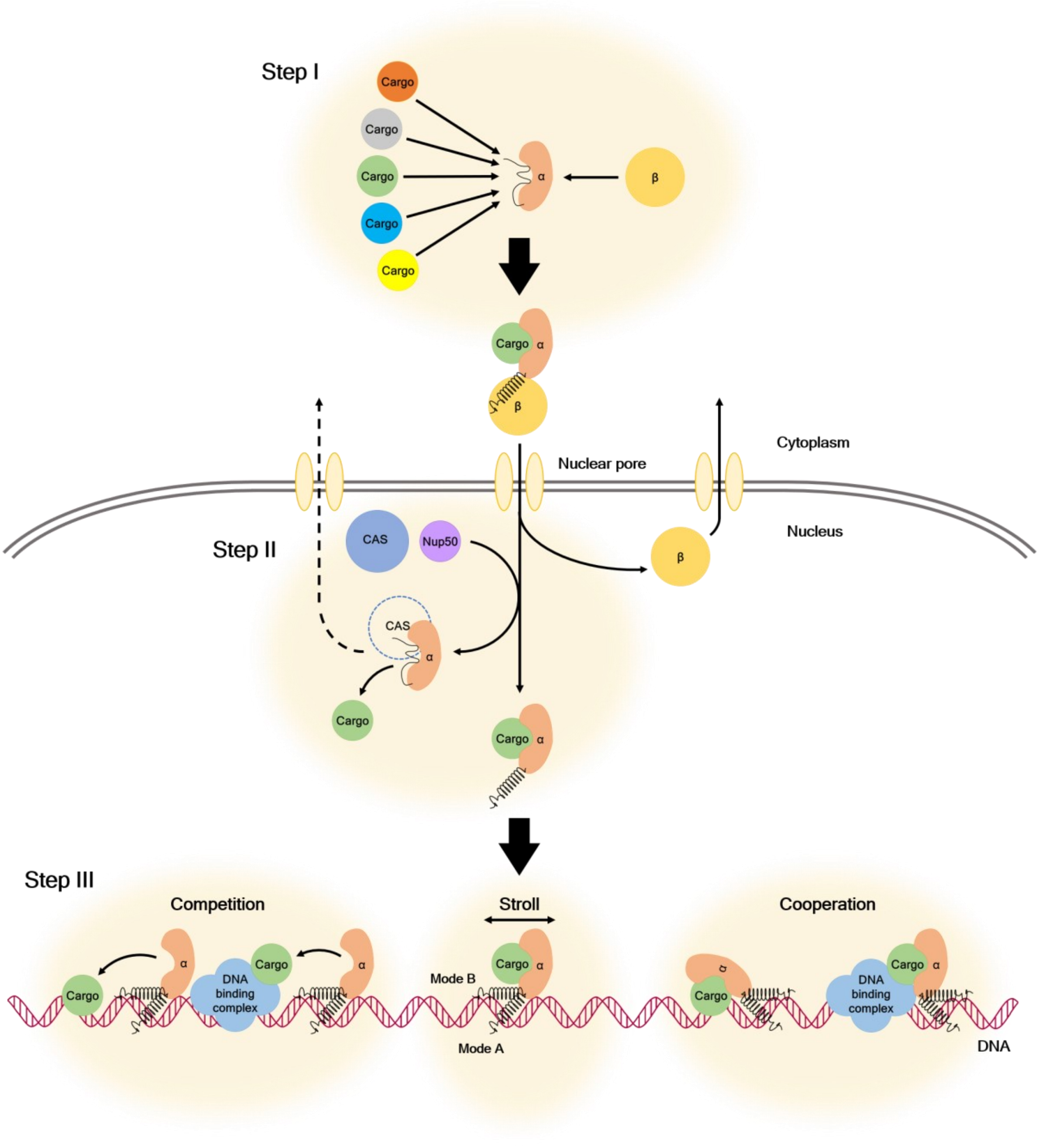


**Figure S11.** Competitive and cooperative steps of importin α–dependent transport and chromatin association involved in cargo destination. The steps are as follows: (I) selections outside the nucleus on the NLS-binding site, by competition between IBB and cargo, and between different cargos, (II) inside the nucleus just after withdrawal of importin β1, cargo is determined whether to be released by competition among IBB, Nup50 (1) or CAS (2, 3) and cargos, and (III) deep inside the nucleus, strolling around DNA, cargo is released by competition of cargo binding between importin a and DNA target sites or other DNA-binding proteins, otherwise the cargos may also cooperatively bind their target sequences with importin α.
