## supplementary methods for "Importin α2 association with chromatin: Direct DNA binding via a novel DNA binding domain"

### Supporting Methods

**1. Supporting methods for Proteins used in gel shift analysis**
**2. Supporting methods for ChIP analyses**
**3. Supporting methods for homology analyses**
**4. Supporting methods for computational analyses**
**5. Supporting methods for the Schiffer-Edmundson helical wheel projection analysis of the NAAT domain**
**6. Supporting methods for NMR**
**7. Supporting methods for molecular phylogenetic analysis coupled with alignment of functional amino acids in IBB**

### 1. Supporting methods for the gel-shift analysis

#### 1.1. Protein purification

Recombinant proteins were purified from pGEX constructs by using a modified version of a previously reported technique (1). The modified protocol has been summarized briefly as follows. Plasmid constructs of proteins were transformed to BL21 (DE3) and were cultured in 200 mL of LB at 30°C for 5 h. Then were cultured with induction of expression by adding 0.1 μM IPTG at 24°C for 6 hours, lysed with 100 μg/mL lysozyme in PBS with PMSF and 0.5% NP40 followed by sonication for 10 s four times with a 10-s interval at 40% maximum amplitude in level 4 with SONIFIER 250 (Branson, US). The supernatants were fractionated by centrifugation at 10000 rpm at 4°C for 10 min. GST-fused proteins were concentrated from the supernatant using GST accept (Nacalai Tesque Inc., Japan.) according to the manufacturer's protocol. The GST tag was cut off using Prescission Protease (GE Life Science, US) or Thrombin (Nacalai Tesque Inc., Japan.) depending on the vector used, in accordance with the manufacturer's protocol. Proteins were purified using PD 10 Desalting Columns (GE Life Science, US) using buffer (20 mM HEPES, 110 mM KOAc, 5 mM NaOAc, 2 mM MgOAc, 1mM EGTA, and 2 mM DTT with protease inhibitors).


### 2. Supporting methods for ChIP analyses

#### 2.1. Quality assessment of ChIP samples

IP-western blot (WB) was performed to check the efficiency of importin α2 immuno-precipitation (IP) using the ChIP protocol. The antibody used for IP was rabbit polyclonal antibody (Abcam, UK), as mentioned in the materials and methods in the text, and the antibody used for WB was goat polyclonal antibody for KPNA2 (Novus Biologicals, US).

### 3. Supporting methods for homology analyses

#### 3.1. Methods for homology analysis with known nucleic acid-binding proteins

Protein BLAST was performed with the following parameters: query sequence, AARLNRFKNKGKDSTEMRRRRISNVELRKAKKDEQMLKRRNVSSF for the whole IBB basic region and KDSTEMRRRRISNVELRKAKKDEQMLKRR for the core short peptide; database: Protein Data Bank (PDB); algorithm, BLASTp (protein-protein BLAST), with automatic adjustment of parameters for short input sequences; expected threshold, 5; word size, 3; matrix, PAM30; gap costs, Existence 8, Extension 1; compositional adjustments: conditioned compositional score matrix adjustments.

Conversely, we extracted all PDB entries (from the database on 25th, September 2019) that included both protein and DNA or RNA molecules at the same time, and we matched them with the output of BLASTp described above. In addition, for the protein sequences with the RRRR motif, each structure was individually examined, and the secondary structure and the interaction between the motif and the nucleic acid were confirmed.

### 4. Supporting methods for computational analyses

#### 4.1. Molecular docking

We constructed the importin α2 IBB domain/DNA complex. The helix structures were extracted from the importin β binding (IBB) domain of importin α in complex with the importin β1 structure (PDB ID: 1QGK) (1). Several amino acids in the extracted helix structures were manually changed to match the sequence of mouse importin α2. For DNA, we generated a canonical B-form DNA structure using the NAB molecular manipulation language (2) in the AmberTools package (3). The first 20-bp sequence of the Oct3/4 upstream region tested in gel-shift assays (upstream-1) was used to construct the DNA model structure.

The helix and DNA structures were docked using AutoDock vina (4). The docking centre was set to the centre of the mass of the DNA and the docking box size was set to [x = 50 Å, y = 50 Å, z = 50 Å] to cover the whole DNA structure. This generated 10 docking poses, and we selected the top scoring docking structure for further molecular dynamics simulation to optimise the docked model structure.

#### 4.2. Molecular dynamics (MD) simulation

The net charge of the simulation system was neutralised by adding counterions ($Na^+$) to the docked model structure with the Leap module in the Amber16 package (3). The docked model structure with counterions was placed at the centre of the water box, and the water molecules were modelled using the TIP3P 3-point charge model potential (5). The size of the water box was chosen such that the distance between every atom in the protein, DNA, and the boundary of the water box was at least 15 Å. The fully solvated system was energy-minimised using 100 steps of the steepest descent, followed by the conjugate gradient method until the energy gradient was less than 0.0001 kcal/mol.

We started with the energy-minimised structure and performed MD simulations to ensure better conformational sampling using the pmemd module in the Amber 16 package (3). Trajectories were calculated for 30 ns. The molecular mechanical force field of *ff*14SB (6) for protein and *OL*15 (7) for DNA was adapted, with the time step of integration set to 1 fs. All bond lengths involving hydrogen atoms were constrained to their respective equilibrium values by the SHAKE method (8). The non-bonded van der Waals interactions were estimated using the Lennard-Jones potential, and the electrostatic energies and forces were calculated using the particle-mesh Ewald summation algorithm (9) with a cut-off distance of 10.0 Å. The system was gradually heated to 300 K during the first 50 ps at a heating rate of 6 K/ps. Subsequently, the temperature and pressure were maintained at a constant 300 K and 1 atm, respectively, by applying the thermostat of Berendsen et al. (10), with a coupling time constant of 1.0 ps. The MD trajectories were sampled every 10 ps and employed for root mean squared deviation (RMSD) calculations, principal component analysis (PCA), and binding free energy estimation using AmberTools 16 (3).

**4.3. Estimation of binding free energies**

Binding free energies between the N-terminal α-helix in the importin α2 IBB domain and DNA ($\Delta G$) were estimated by the molecular mechanics generalised born surface area (MM-GBSA) implicit solvent model (11). The $\Delta G_{\mathrm{MMGBSA}}$ were calculated every 100 ps, from 1 ns to 30 ns for the wild type and from 1 ns to 10 ns for variants. The $\Delta G_{\mathrm{MMGBSA}}$ at each time, $t$ ps, was calculated from 100 snapshots saved every 10 ps from ($t$-990) ps to $t$ ps. All the $\Delta G_{\mathrm{MMGBSA}}$ calculations were based on the GB-OBC model (12) and interior and exterior dielectric constants of $\varepsilon_{\mathrm{in}} = 1$ and $\varepsilon_{\mathrm{out}} = 80$, respectively.

The implicit solvent GBSA model accounts for the solvent entropic effect but not the configurational entropic effect of the solute. The configurational entropies of the solute were calculated using PDB2ENTROPY and PDB2TRENT (13). For the calculations, the MD trajectories were as same as those described for $\Delta G_{\mathrm{MMGBSA}}$.

### 5. Supporting methods for the helical wheel projection

#### 5.1. Methods for the Schiffer-Edmundson helical wheel projection analysis of the NAAT domain

The helical wheel (1) was created using NetWheels: Peptides Helical Wheel and Net projections maker (2).

1. Schiffer, M. and Edmundson, A. B. (1967). Use of helical wheels to represent the structures of proteins and to identify segments with helical potential. *Biophysical Journa, 7, 121-35.*
2. Mól, A. R, Castro, M, S. Fontes, W. NetWheels: A web application to create high quality peptide helical wheel and net projections. bioRxiv 416347; doi: https://doi.org/10.1101/416347 (preprint)

### 6. Supporting methods for NMR

#### 6.1. Equilibrium analysis to determine the dissociation constant and stoichiometry

The results of NMR titration experiments of 15-bp DNAs with a 30-aa long peptide (importin α2_21-50, the core part of the arginine-rich region in mouse importin α2) were analysed to estimate the binding strength and stoichiometry. At first, the imino proton region of each 1D NMR spectrum at each peptide concentration in the titration experiments was decomposed into twelve peaks by non-linear fitting using Lorentz peak functions (1, 2),

$$\mathrm{f(x)}_i = 2\mathrm{h}_i\mathrm{w}_i/(4\pi(\mathrm{x}-\mathrm{u}_i)^2 + w_i{}^2) + \mathrm{b}_i$$

where $\mathrm{h}_i$, $w_i$, $\mathrm{u}_i$, and $\mathrm{b}_i$ are parameters for peak height, peak width, peak centre, and baseline for $i$-th peak respectively.

Figure S7 shows the representative results of the decomposition of the spectra for SOX-POU DNA and for random-sequence DNA in the absence of the peptide. The decomposed peaks are summarized in Table S2 and two examples of the changes of the decomposed peaks are shown in Figure 4G and H for SOX-POU DNA and random-sequence DNA, respectively. The observed spectral change obviously showed two phases, and the findings indicated that the binding of the peptide to DNA had at least two modes that probably correspond to three equilibrium states from free DNA, DNA-peptide complex-1, and DNA-peptide complex-2. For the SOX-POU core sequence, the association for the first step and for the second step seemed to be saturated at peptide concentrations of 100 μM and 200 μM to 250μM, respectively, against the 50 mM DNA duplex. Therefore, we assumed a two-step sequential binding process in which the stoichiometry is 1:2 (DNA:peptide) for the first step and 1:3 (DNA:peptide) for the second step. On the other hand, for the random-sequence DNA, the association for the first step and for the second step seemed to be saturated at peptide concentrations of 150 μM and 200 μM, respectively, against the 50 mM DNA duplex. Although the binding seemed to be relatively weak and the saturation points were ambiguous, we assumed a two-step sequential binding process in which the stoichiometry is 1:3 (DNA:peptide) for the first step and 1:1 (DNA:peptide) for the second step.

Under the assumption that the two processes are independent and completely sequential, the changes in the chemical shift and peak height were then analysed by non-linear least-square fitting to estimate the binding strengths and stoichiometries.

Assuming a model described above, the chemical shift change of the *i* th peak $\Delta\delta_{obs}(i)$ is expressed as follows,

$$\Delta\delta_{obs}(i) = \Delta\delta1_{obs}(i) + \Delta\delta2_{obs}(i)$$

where $\Delta\delta1_{obs}(i)$ is chemical shift change in the first step and $\Delta\delta2_{obs}(i)$ is that in the second step. Each component of the total chemical shift change ( $\Delta\delta1_{obs}(i)$ and $\Delta\delta2_{obs}(i)$ ) can be expressed as a product of the chemical shift change of peak *i* at saturation point and the population of the bound form of binding sites on the DNA (3),

$$\Delta\delta1_{obs}(i) = \Delta\delta1_{sat}(i)\frac{[p]_t - [p]}{[D_{site1}]_t}$$

$$= \frac{\Delta\delta1_{sat}(i)}{[D_{site1}]_t} \cdot \frac{([p]_t + K_{d1} + [D_{site1}]_t) - \sqrt{([p]_t + K_{d1} + [D_{site1}]_t)^2 + 4K_{d1}[p]_t}}{2}$$

where $\Delta\delta1_{sat}(i)$, $[p]_t$, $[p]$, $[D_{site1}]_t$, and $K_{d1}$ are the chemical shift change of peak *i* at the saturation point, the total concentration of the peptide, the concentration of unbound free peptide, and the total concentration of the binding sites on DNA for the first step.
After the saturation of the first step, the concentration of free peptide is approximately written as follows,

$$[p] = [p]_t - [D_{site1}]_t$$

Then for the chemical shift change of the peak, *i* for the second step is

$$\Delta\delta2_{obs}(i) = \Delta\delta2_{sat}(i)\frac{[p]_t - [p]}{[D_{site1}]_t}$$

$$= \frac{\Delta\delta2_{sat}(i)}{[D_{site2}]_t} \cdot \frac{([p]_t + K_{d2} + [D_{site2}]_t) - \sqrt{([p]_t + K_{d2} + [D_{site2}]_t)^2 + 4K_{d2}[p]_t}}{2}$$

where $\Delta\delta2_{sat}(i)$, $[D_{site2}]_t$, and $K_{d2}$ are the chemical shift change of peak *i* at saturation point, the total concentration of the binding sites on DNA, and the dissociation constant for the second step.


1. Cavanagh, J., Fairbrother, W. J., Palmer III, A. G. and Skelton, N. J. (1996) Protein NMR spectroscopy principles and practice, Academic Press, San Diego, CA.
2. Tauler, R. (1995). *Chemometr. Intell. Lab. Syst.*, **30**, 133
3. Inase, A., Kodama, T.S., Sharif, J., Xu, Y., Ayame, H., Sugiyama, H., and Iwai, S. (2004) Binding of distamycin A to UV-damaged DNA. *J Am Chem Soc,* **126**, 11017-11023


### 7. Supporting methods for molecular phylogenetic analysis coupled with alignment of functional amino acids in IBB

#### 7.1. Methods

##### 7.1.1. Relative solvent accessible area of each amino acid side chain

Solvent-accessible surface area (SASA) of each amino acid side chain was calculated using a home developed program written in c language. The program has been derived from STRIDE (1) so as to evaluate amino acid main chain and side chain separately. The relative solvent accessible area (RAS) was calculated as a relative ratio of the solvation area of each amino acid to the theoretical maximum value for the amino acid type calculated from average coordinates of peptide sequence GGXGG where X was each amino acid type. The coordinates of PDB:1QGK chain B (importin $\alpha$ 2 IBB domain) were used for the calculation of RAS of IBB domain side chains.

### 7. Supporting methods for molecular phylogenetic analysis coupled with alignment of functional amino acids in IBB

#### 7.1. Methods

##### 7.1.1. Relative solvent-accessible area of each amino acid side chain

Solvent-accessible surface area (SASA) of each amino acid side chain was calculated using a home developed program written in c language. The program has been derived from STRIDE (1) so as to evaluate the amino acid main chain and side chain separately. The relative solvent-accessible area (RAS) was calculated as a relative ratio of the solvation area of each amino acid to the theoretical maximum value for the amino acid type calculated from the average coordinates of the peptide sequence GGXGG, where X was each amino acid type. The coordinates of PDB:1QGK chain B (importin $\alpha$ 2 IBB domain) were used for the calculation of RAS of IBB domain side chains.

#### 7.1.2. Evolutionary conservation of each residue

The degree of conservation of each residue was determined using Consurf server (2). Parameters for the homolog search were chosen as follows: homolog search algorithm: HMMER, number of iterations: 1, E-value cut-off: 0.01, protein database: UNIREF90, calculation method: Bayesian, evolutionary substitution model: best model.

The sequence logo of IBB (residues from 11 to 54, Figure 2E) was created by WebLogo server (Crooks GE, Hon G, Chandonia JM, Brenner SE (2004) WebLogo: A sequence logo generator, Genome Research, 14:1188-1190) for the multiple sequence alignment containing 250 IBB sequences that was obtained from the above analysis.

#### 7.2. Interpretation

The multifunctionality of the IBB domain was evaluated by molecular phylogenetic analysis of the IBB domain. The degree of conservation of each residue in the amino acid sequence of the IBB domain was examined, after which the contribution to binding at the complex interface with importin β1 was considered. As expected, the degree of conservation of the amino acids involved in the binding interface was high. This was consistent with a mutation rate that is approximately proportional to the relative solvent accessibility (RSA, a measure of residue "buriedness") of the amino acid residues, and highly buried residues show a high degree of evolutionary conservation (3-7). Conversely, in addition to the residues buried by binding to importin β1, multiple highly conserved residues were detected in the IBB domain of importin α, strongly suggesting that the importin α IBB domain in helix structures has an important function in addition to binding with importin β1.
