## supplementary discussions for "Importin α2 association with chromatin: Direct DNA binding via a novel DNA binding domain"

**Results and Discussion of MD simulation**

**1. Overall structure**

We tested the stability and equilibrated the initial model structure obtained by molecular docking by a 30-ns MD simulation. The initial α-helix position was improved, yielding a root-mean-square (RMS) deviation of 8 Å for the α-helix position from the initial model averaged over backbone heavy atoms (black line in Figure S5A). Most of this positional adjustment occurred in the early phase of the equilibration (~4 ns), and very little additional drift was observed in the α-helix position during 4–21-ns period (black line in Figure S5A), with the remaining change of ±1.3 Å in the RMS deviation. After 21 ns, the α-helix position showed a large drift and reached a second stable state at 25–30 ns, with RMS deviation of 18 ± 1.3 Å (black line in Figure S5A). No structural changes were observed in the DNA and α-helix itself throughout the simulation (red and blue lines in Figure. S5A). These results indicate two binding modes for the α-helix to DNA in our simulation, mode A and B. In the process of the binding mode change from mode A to mode B, the α-helix transits on the DNA (Figure S5B).

**2. Binding free energy analysis**

We investigated the binding mechanism in greater depth by estimating the binding free energy between the α-helix and DNA. The binding free energies fluctuated during a period of each binding mode (Figure S5C), at -526.05 ± 36.28 kJ/mol for mode A and -602.96 ± 27.91 kJ/mol for mode B. A snapshot showing the lowest free energy in each binding mode was chosen as a representative structure of each mode.

Our binding free energies, estimated using the MMGBSA method, were relative rather than absolute values. We assessed the strength of the wild-type α-helix binding to DNA by also evaluating the binding affinities of three variants of the α-helix, 28A4, 39A5, and 49A3, which were examined experimentally. The mutations (28RRRR → 28AAAA, 39RKAKK → 39AAAAA and 49KRR → 49AAA) were introduced manually in the representative structure for both modes A and B, and then six 10-ns MD simulations were performed, starting from these initial structures (three variants, 28A4, 39A5, and 49A3, for both modes A and B).

The binding free energies for the three variants of the α-helix that showed no binding to DNA experimentally were -509 to -296 kJ /mol (Figure S5D, and E). The DNA binding of these α-helix variants were weakened in the order of 28A4 ≈ 39A5 > 49A3 for both modes A and B, and the tendency was consistent with our experimental data. A comparison of the binding free energies between the wild type and the three variants strongly suggested that the α-helix in both modes A and B bound to the DNA. Despite this binding to DNA, the binding affinity of the α-helix to DNA would not be strong because we noted transition of the α-helix on the DNA from mode A to mode B in our short-timescale MD simulation. These results led us to assume the that α-helix (importin α2) binding to DNA is neither strong nor weak; therefore, the α-helix (importin α2) would transit on the DNA.

**3. Free energy landscape**

The stochastic dominant protein/DNA complex structure must have a minimal free energy conformation. This configuration can be explored by constructing a free energy landscape (FEL) from the MD trajectory. In the present study, free energies were mapped onto the configurational space formed by the principal components (PCs). We elucidated the PCs of the α-helix/DNA complex configurations by subjecting the MD trajectory to PCA. This method projects the dynamics captured in the trajectory onto a reduced dimensionality. In the PCA, a covariance matrix was constructed from the MD trajectory and then diagonalised to obtain the solution of eigenvectors and eigenvalues for the matrix. Here, we extracted the first two PCs. The cumulative contribution of these two PCs to the overall dynamics was 64%.

Both A and B modes for α-helix/DNA binding resulted in configurational clusters with lower free energies in FEL (Figure S5F). This indicated small configurational variations of both A and B modes by restricting the relative motions between the α-helix and DNA. Energy barriers were present between A and B models on FEL; however, these barriers would be low and easily overcome by thermal fluctuations. This is supported by the fact that we directly observed the binding mode change from A to B along with the transition of the α-helix on the DNA during our MD simulation. The FEL analysis allowed us to conclude that the binding mode of the α-helix (importin α2)/DNA is elusive; this is consistent with the nature of the non-specific protein-DNA complex previously proposed by many experimental and theoretical studies (see review article by (1) and references therein).
